# Nanoparticles use magnetoelectricity to target and eradicate cancer cells

**DOI:** 10.1101/2024.10.13.618075

**Authors:** John Michael Bryant, Emmanuel Stimphil, Victoria Andre, Max Shotbolt, Elric Zhang, Veronica Estrella, Kazim Husain, Joseph Weygand, Doug Marchion, Alex Sebastian Lopez, Dominique Abrahams, Shawnus Chen, Mostafa Abdel-Mottaleb, Skye Conlan, Ibrahim Oraiqat, Vaseem Khatri, Jose Alejandro Guevara, Shari Pilon-Thomas, Gage Redler, Kujtim Latifi, Natarajan Raghunand, Kosj Yamoah, Sarah Hoffe, James Costello, Jessica M. Frakes, Ping Liang, Sakhrat Khizroev, Robert A. Gatenby, Mokenge Malafa

## Abstract

This study presents the first in vivo and in vitro evidence of an externally controlled, predictive, MRI-based nanotheranostic agent capable of cancer cell specific targeting and killing via irreversible electroporation (IRE) in solid tumors. The rectangular-prism-shaped magnetoelectric nanoparticle is a smart nanoparticle that produces a local electric field in response to an externally applied magnetic field. When externally activated, MENPs are preferentially attracted to the highly conductive cancer cell membranes, which occurs in cancer cells because of dysregulated ion flux across their membranes. In a pancreatic adenocarcinoma murine model, MENPs activated by external magnetic fields during magnetic resonance imaging (MRI) resulted in a mean three-fold tumor volume reduction (62.3% vs 188.7%; *P* < .001) from a single treatment. In a longitudinal confirmatory study, 35% of mice treated with activated MENPs achieved a durable complete response for 14 weeks after one treatment. The degree of tumor volume reduction correlated with a decrease in MRI T_2_* relaxation time (*r* = .351; *P* = .039) which suggests that MENPs have a potential to serve as a predictive nanotheranostic agent at time of treatment. There were no discernable toxicities associated with MENPs at any timepoint or on histopathological analysis of major organs. MENPs are a noninvasive alternative modality for the treatment of cancer.

**Summary:** We investigated the theranostic capabilities of magnetoelectric nanoparticles (MENPs) combined with MRI via a murine model of pancreatic adenocarcinoma. MENPs leverage the magnetoelectric effect to convert an applied magnetic field into local electric fields, which can induce irreversible electroporation of tumor cell membranes when activated by MRI. Additionally, MENPs modulate MRI relaxivity, which can be used to predict the degree of tumor ablation. Through a pilot study (n=21) and a confirmatory study (n=27), we demonstrated that, ≥300 µg of MRI-activated MENPs significantly reduced tumor volumes, averaging a three-fold decrease as compared to controls. Furthermore, there was a direct correlation between the reduction in tumor T_2_ relaxation times and tumor volume reduction, highlighting the predictive prognostic value of MENPs. Six of 17 mice in the confirmatory study’s experimental arms achieved a durable complete response, showcasing the potential for durable treatment outcomes. Importantly, the administration of MENPs was not associated with any evident toxicities. This study presents the first in vivo evidence of an externally controlled, MRI-based, theranostic agent that effectively targets and treats solid tumors via irreversible electroporation while sparing normal tissues, offering a new and promising approach to cancer therapy.

## Introduction

Locally advanced pancreatic adenocarcinoma (LAPC) is a very morbid malignancy, with few therapeutic options and a limited life expectancy of 6 months to 1 year^1^. It is projected to become the second leading cause of cancer-related death in the United States by 2030^2^. To overcome the limitations of existing modalities and meet the urgent need for new and effective therapies for LAPC and other treatment-resistant cancers, this study aims to evaluate the potential use of magnetoelectric nanoparticles (MENPs) as a new therapeutic agent^3^.

Nanoparticles (NPs) are nanoscale structures (1-100 nm) with properties that can be modified for various biomedical applications. The customizability of nanoparticles enables versatile and unique mechanisms. A new smart NP, defined as those that change their function in response to certain external stimuli, that holds great potential for cancer care is the MENP^4^. MENPs are defined by their unique magnetoelectric (ME) effect, which arises from their core-shell interactions (eg, CoFe_2_O_4_@BaTiO_3_) and allows MENPs to efficiently convert magnetic fields into local electric fields and vice-versa^5^ (**Fig. 1a**). With this ME effect, MENPs can deliver drug payloads with high active external control and generate local electric fields to activate neurons^6–14^. A unique consequence of the ME effect is that MENPs can specifically target cancer on a cellular level due to the non-polar and highly conductive cell membranes of cancer cells^15–17^. Given their cancer targeting capabilities and ability to generate large local electric fields, MENPs may be able to induce a targeted, locally ablative cell-kill ability via irreversible electroporation (IRE)^18,19^; however, this has not previously been demonstrated.

**Fig. 1:**
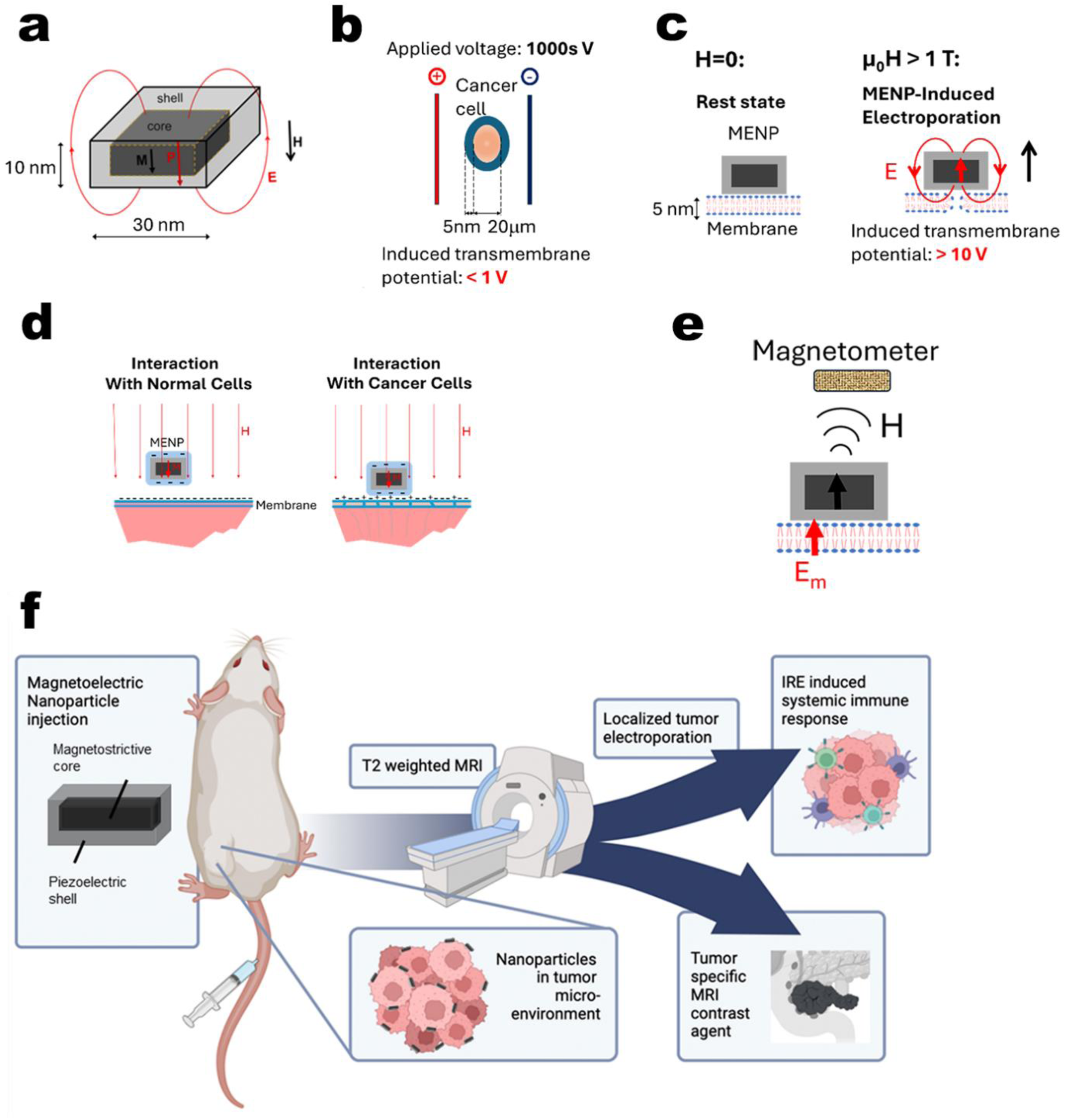
MENPs’ underlying physics and local field generation for enabling IRE and MRI detection. **a MENP core-shell structure and correlated electric and magnetic field.** Core-shell MENPs are made of lattice-matched magnetostrictive cores and piezoelectric shells, and they exhibit an ME effect because of strain propagation through the surface interface. An applied magnetic field, H, induces a magnetization, M, that in turn leads to an electric dipole field, E, applied to the membrane. m_0_ (=4π × 10^-7^ H/m) is the magnetic constant. **b Conventional IRE.** For comparison, in the conventional IRE setting, application of a 1 kV voltage induces a transmembrane voltage of <1 V because of the conductive nature of TME. **c MENP-controlled targeted IRE.** If no magnetic field is applied, the MENP is at rest and only weakly polarized. In a conservative estimate, MENPs are capable of generating a transmembrane voltage of 10 V (ie, an order of magnitude greater than conventional IRE). **d Interaction with normal cells and cancer cells.** MENPs prefer to target cancer cells because cancer cells have more depolarized membranes than normal cells. When activated by a small external magnetic field, there is a greater probability for MENPs to overcome repulsive electrostatic forces, thus reaching the regime of the attractive van der Waals force, with the nonpolar and more conductive cancer cellular membrane than with normal cellular membranes. The nanoparticles display a negative zeta potential in the TME. **e MRI relaxivity modulation.** The MENP’s ferrimagnetic core increases the dephasing of proton spins because of their inhomogeneous local magnetic fields. **f Theranostic application of MENPs.**

IRE is an effective therapy for LAPC that can ablate the tumor microenvironment (TME) and induce an immunostimulatory effect^20^. IRE relies on the generation of sufficient electric fields across the membrane that permanently rearrange the membrane’s lipid molecules by creating toroid-shaped pores^21^. The reversibility of this process depends on the type, size, shape, and membrane bilayer composition of the cell as well as the strength and frequency of the applied electric field. With conventional IRE, physical electrodes are inserted into a cancer site and large electric fields (100 kV/m) are generated between them to achieve a transmembrane voltage of <1 V^22^. Regardless of whether it is performed percutaneously or surgically, the invasive nature of conventional IRE limits its use^21^. Additionally, because the TME is conductive, application of thousands of volts between the electrodes can also cause significant electric currents to flow through nearby normal tissues (**Fig. 1b**). Hence, conventional IRE is considerably toxic to sensitive nearby normal tissue^23–25^. In contrast, MENPs may be able to overcome the limitations of conventional IRE by eliminating both the need for an invasive procedure and for large electric fields with high currents.

MENPs are theoretically capable of delivering ablative IRE in a highly controlled, targeted, and localized manner (ie, nanometers within their immediate proximity) while simultaneously sparing normal cells (**Fig. 1c**) because of their inherent ME effect, size and shape, and cellular membrane electrostatic forces^15,18,26^. MENP’s electric field can generate a significantly higher transmembrane potential (>10 V) on the cellular membrane than that created with conventional IRE (<1 V) without causing its associated side effects. Furthermore, MENPs display high-specificity targeting capabilities after intravenous administration, as previously demonstrated through an in vivo murine model^16^.

Because of their ferrimagnetic cores, MENPs can target a known tumor location via application of an external magnetic field^22^. There are 2 counterbalancing forces acting on nanoparticles when they approach the cellular membrane: the attractive van der Waals force and the repulsive electrostatic force. For both normal cells and cancer cells, the van der Waals force is equally proportional to the interface surface area; thus, they prefer to orient the nanoparticles so that the nanoparticle’s largest surface is parallel to the membrane surface, regardless of the cell type. In contrast, the electrostatic force depends on the cell type. Given that the zeta potential of MENPs is negative in the TME, MENPs exhibit a preference towards aggregating upon the more depolarized and conductive cancer cell membranes^15,16^ (**Fig. 1d**). The physics of the electrostatic force depends on the magnetic field application and the interplay between the field-controlled surface charge and dipole moment, as has been previously described in detail^27^.

Additionally, MENPs can also serve as T_2_ and T_2*_ signal modulators on MRI^28–30^ (**Fig. 1e**). In conjunction with strong local electric fields, the non-zero magnetic fields that arise due to MENPs’ ferrimagnetic cobalt-iron oxide core (CoFe_2_O_4_) cause local magnetic field inhomogeneities, thus decreasing spin-spin relaxation times in a concentration-dependent fashion^31,32^. Hence, we hypothesize that MENPs can serve as a theranostic anti-cancer nanotherapeutics that can both (i) generate externally controlled and targeted ablation via IRE with high specificity towards cancer cells and (ii) simultaneously induce a detectable MRI signal on T_2_/T_2*_ relaxometry within the targeted tumor (**Fig. 1f**).

In this study, we conducted both in vitro and in vivo experiments in conjunction with MRI to elucidate the anti-tumor therapeutic mechanism of action, efficacy, and T T * MRI relaxivity modulating effects of MENPs. We aimed to demonstrate that, when exposed to MRI magnetic fields, MENPs could perform tumor-targeted predictive theragnosis in a pancreatic adenocarcinoma tumor model.

## RESULTS

### Outline of experiments

To test the theranostic potential of MENPs, 2 murine experiments were performed in succession using a flank tumor model to evaluate the in vivo tumor-cell kill effects of MRI-activated MENPs. MRI activation and relaxivity experiments were performed on a 7 T Biospec MRI Scanner (Bruker, Billerica, MA). Both the pilot study and confirmatory study followed similar protocols with minimal changes (**Fig. 2a**). The primary aims of this pilot study were (i) signal modulation of T_2_ MRI mapping studies at M1 (defined as 2 days after baseline) from baseline (M0) and (ii) tumor volume change at M2 (defined as 7 days after baseline) from M0. The mice were immediately euthanized after their M2 MRI scan. The follow-up confirmatory study cohort design was informed by the pilot experiment. The primary aims for the confirmation study were the same, with the addition of characterizing tumor volume response over time and histological analysis of organs to evaluate the effect of MENPs within vital bodily organs.

**Fig. 2:**
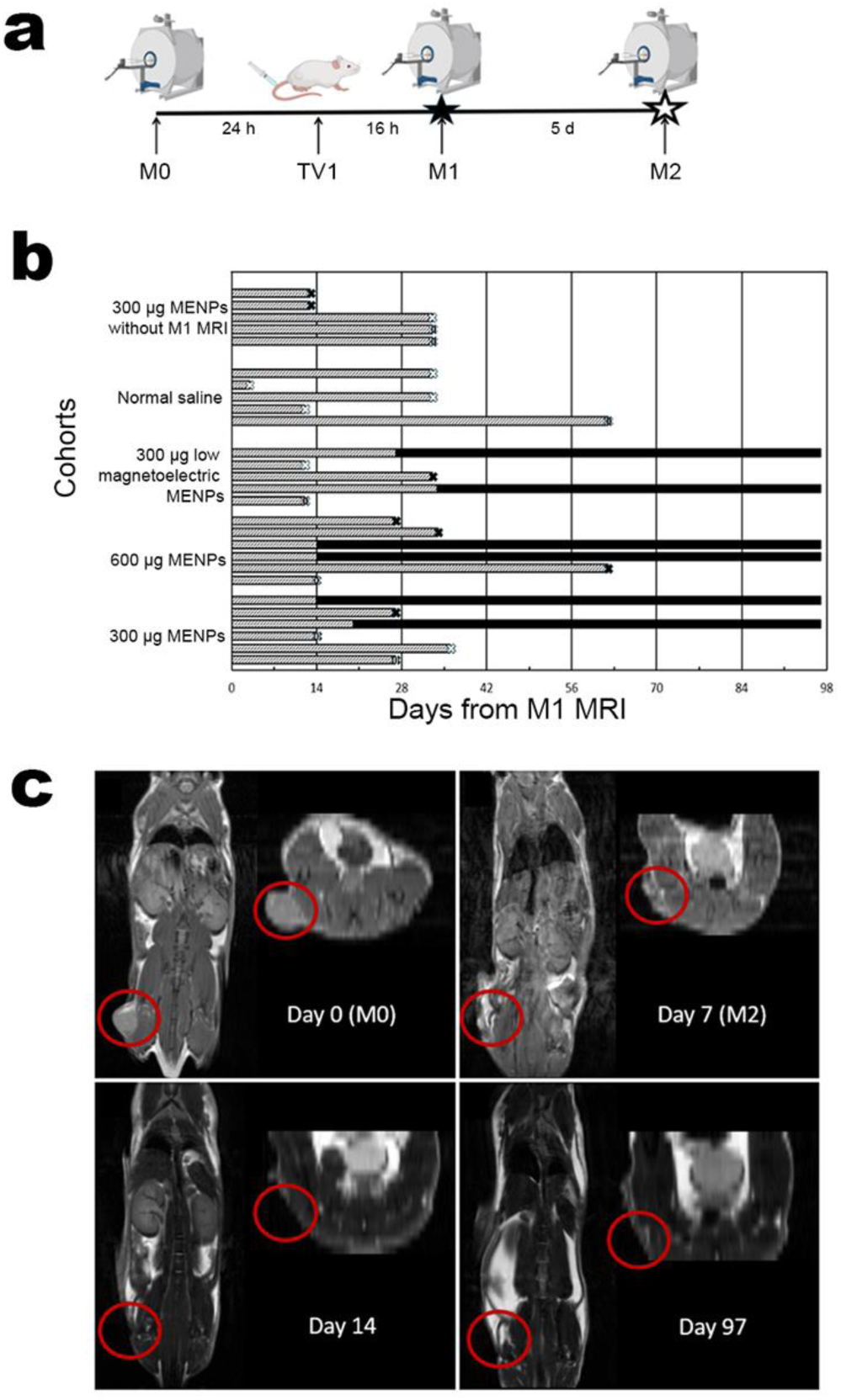
Study schema and time-to-event outcomes. **a Schema for both pilot and confirmatory studies**. The in vivo schema depicts the order and time intervals of the pilot and confirmatory study between the baseline MRIs (M0) and the M2 MRIs. After their tumors grew to sufficient sizes (detailed within each study method), the mice were randomized into cohorts. All mice then underwent a baseline MRI. The next day, all mice received an injection according to their cohort, which consisted of either MENPs of various concentrations in 300 µl of deionized water with MENPs or normal saline without MENPs. During this time, they also had a neodymium magnet placed directly over their flank tumors. These magnets were removed immediately before the M1 MRIs, which served as the endpoint for the relaxivity aim. No further injections, targeting, or imaging were performed for 5 days, following which the M2 MRIs, which served as the endpoint for the tumor volume aim, were performed. In the pilot study, mice were then euthanized. In the confirmatory study, mice were observed for 12 weeks (14 weeks from baseline). In this figure, M0 indicates baseline MRI, TV1 indicates tail vein injection, M1 indicates relaxivity aim MRI, and M2 indicates tumor volume aim MRI. **b Time-to-event waterfall plot of confirmatory study mice from timepoint M1 (ie, MENP activation via MRI).** Two mice within the 300, 600, and LM cohorts (ie, 6 mice in total) achieved a CR (as determined via exam and MRI). After achieving a CR, none of the mice experienced any evidence of disease recurrence. **c** Example mouse from 300 cohort in the confirmatory study that achieved a durable CR. The top left image shows a coronal (left) and axial (top right) view of the T_1_-weighted (from T_1_ relaxometry mapping study) MRI at baseline, showing a 107.9 mm^3^ flank tumor (circled in red). The top right image shows a coronal and axial view of tumor (circled in red) at M2 (day 7 from baseline), demonstrating a significant decrease in tumor volume to 32.3 mm^3^ (31.8% of baseline). The bottom left image shows a coronal and axial view of T_2_-weighted (from RARE MRI [turbo spin echo rapid acquisition with relaxation enhancement]) MRI, showing a CR (circled in red) at day 14 from M0 (one week from M2). The bottom right image shows a coronal and axial view of a durable CR (circled in red) after 14 weeks.

### Pilot in vivo murine study

For the pilot experiment, 30 (15 female and 15 male) 6- to 8-week-old immunocompetent C57BL/6 mice (Charles River Laboratories, Wilmington, MA) were obtained and allowed to acclimate to the vivarium for 1 week before being subcutaneously inoculated with right flank tumors of 1,000,000 KPC961 pancreatic adenocarcinoma cells. The 10 female and 11 male mice with the largest tumors, as measured with calipers 3 weeks after inoculation, were randomized into 5 cohorts based on treatment: 300 µg of MENPs (ie, the 300 cohort; n=5), 600 µg of MENPs (ie, the 600 cohort; n=5), 60 µg of MENPs (ie, the 60 cohort; n=5), 300 µg of low–ME effect MENPs (ie, the LM cohort; n=3), and normal saline (ie, the NS cohort, which acted as the MENPs causal effect control; n=3). According to the calibrated dye test (described in Methods), the LM cohort’s dose showed approximately half of the ME effect as the normal MENPs. The median tumor size across all cohorts at M0 was 83.0 mm^3^ (interquartile range [IQR], 41.2-119.5 mm^3^; *F*_4,16_ = 2.323; *P* = .101), and the median and mouse weight was 25.3 g (IQR, 20.9-26.5 g; *F*_4,16_ = .982; *P* = .445).

All cohorts underwent MRI mapping studies at timepoint M0 on a small animal 7 T MRI that consisted of T_2_*, T_2_, and T_1_ maps. One day after the M0 MRIs, the mice underwent tail vein treatment injections (**Extended Data Fig. S1a**) that corresponded with their cohort in a solution volume of 300 µL. Immediately after tail vein injection, every mouse had a neodymium magnet (**Extended Data Fig. S1b**) placed over their tumors that generated approximately 0.15 T over the tumor. The magnet was placed for approximately 16 hours overnight using VetBond Tissue Adhesive (3M, Saint Paul, MN) to promote accumulation of the MENPs at the targeted site. After magnet removal, all mice underwent repeat MRI mapping studies (timepoint M1). The female mice were rehoused together after completion of M1, but the male mice could not be rehoused together because of home-cage aggression and were housed separately for the remainder of the experiment. MRI mapping was repeated five days after M1 (ie, timepoint M2). All mice were euthanized immediately after M2 MRIs.

Tumor volume and relaxivity data for the pilot study are summarized in **Extended Data Table S1**. The ANOVA analyses of absolute T_2_ relaxation changes from baseline yielded *F*_4,16_ = 3.284 (*P* = .038), and relative T_2_ relaxation yielded *F*_4,16_ = 3.939 (*P* = .21). The ANOVA analyses of T_2_* absolute relaxation changes from baseline yielded *F*_4,16_ = 13.815 (*P* < .001), and T_2_* relative relaxation yielded *F*_4,16_ = 18.147 (*P* < .001). We observed statistically significant absolute changes in tumor volume (*F*_4,16_ = 7.882; *P* < .001) and a trend toward statistically significant relative changes (*F*_4,16_ = 2.932; *P* = .054).

Least significant difference (LSD) analyses for the pilot study are summarized in **Extended Data Table S2**. The 300 and 600 cohorts demonstrated the greatest relative tumor shrinkage, with reductions of 72.3% and 49.8% from baseline (ie, M2/M0), respectively. In contrast, the NS cohort experienced an increase of 265%. Compared to NS cohort, the mean differences in tumor shrinkage were –157.9% (*P* < .001) for the 300 cohort and –175.8% (*P* < .001) for the 600 cohort. Additionally, the drop in T_2_* relaxation times from M1 to baseline was 61.4% for the 300 cohort and 59.1% for the 600 cohort, compared to an increase of 151.7% for the NS cohort. Compared to NS controls, the mean differences in T_2_* relaxation times were –84.0% (*P* < .001) for the 300 cohort and –83.7% (*P* < .001) for the 600 cohort. There were no differences in tumor volume response based on sex (mean difference, 2.2%; *t*(19) = .056; *P* = .956), and no mice lost more than 7% of their total body weight (**Extended Data Table S1**). There were no statistically significant differences in weight changes between cohorts from M0 to M2 (*F*_4,16_ = 2.318; *P* = .102).

### Confirmatory in vivo murine study

To confirm the results from the pilot experiment and assess therapeutic response over a 90-day period after a single dose of MENPs, a larger confirmatory study was performed. The 60 cohort from the pilot study was eliminated, as neither therapeutic nor contrast effects were demonstrated. This cohort was replaced with a new cohort that did not undergo MRI at timepoint M1 (ie, the NM cohort). The NM cohort was added to control for any therapeutic effect the MENP particles may have upon the tumor with only the neodymium magnet. Additionally, the NM cohort did not undergo any relaxivity analyses but instead had T_2_-weighted scans at M0 and M2. Given the longer study duration, 35 female mice were used so that they could be re-housed together after separation during targeting. Additionally, we extended the tumor growth time to 5 weeks before the baseline MRI so that we could test the MENPs’ effect on larger tumors.

The 27 mice with the largest tumors, as measured on T_2_-weighted MRIs 4 weeks after inoculation, were randomized into 5 cohorts based on treatment: 300 µg of MENP (ie, the 300 cohort; n=6), 600 µg of MENP (ie, 600 cohort; n=6), 300 µg of low-ME effect MENP (ie, the LM cohort, with an ME effect of ∼ 50%; n=5), 300 µg of MENP without M1 MRI (ie, the NM cohort, which acted as an MRI field control; n=5), and normal saline (ie, the NS cohort, which acted as a MENP causal effect control; n=5). The NM cohort underwent T_2_-weighted MRIs at timepoints M0 and M2 but not at M1. The median tumor size across all cohorts at time M0 (ie, 5 weeks after inoculation) was 109.6 mm^3^ (IQR, 42.0-235.9 mm^3^; *F*_4,22_ = .110; *P* = .978), and median mouse weight was 20.3 g (IQR, 19.9-20.8 g; *F*_4,22_ = .578; *P* = .682). Mice in the confirmatory study underwent the same schema (**Fig. 2a**) as the pilot study, except for the mice within the NM cohort (as described above).

Tumor volume and relaxivity data for the confirmatory study are summarized in **Extended Data Table S3**. One mouse in the NS cohort met the predefined tumor growth endpoint and was subsequently euthanized before timepoint M2. No mice within the 300 or 600 cohorts demonstrated any tumor growth at M2. The ANOVA analyses for the absolute T_2_ relaxation changes from baseline yielded *F*_3,18_ = 1.837 (*P* = .177), and relative T_2_ relaxation yielded *F*_3,18_ = 1.981 (*P* = .153). The ANOVA analyses for T_2_* absolute relaxation changes from baseline yielded *F*_3,18_ = 3.719 (*P* = .031), and T_2_* relative relaxation yielded *F*_3,18_ = 5.237 (*P* = .009). The tumor volume absolute change from baseline yielded *F*_4,21_ = 3.800 (*P* = .018), and tumor volume relative change yielded *F*_4,21_ = 13.411 (*P* < .001).

LSD analyses for the confirmatory study are summarized in **Extended Data Table S4**. The 300 and 600 cohorts demonstrated the greatest relative tumor shrinkage, with mean reductions of 56.7% and 73.7% from baseline, respectively. In contrast, the NS cohort experienced an increase of 159.4%. Compared to the NS cohort, the mean differences in tumor shrinkage were –115.9% (*P* < .001) for the 300 cohort and –106.2% (*P* < .001) for the 600 cohort. Neither the 300 nor the 600 cohort demonstrated a significant relative change in T_2_* relaxivity from baseline (*P* > 0.05); however, they both demonstrated a significant absolute change of -2.5 ms (*P* = .034) and -2.7 ms (*P* = 0.27), respectively. The NM cohort did not demonstrate a significant change in tumor volume compared to the NS control group (163.4% vs. 159.4%; *P* = .427).

In contrast to the pilot experiment, mice in the confirmatory study were not euthanized at M2. Instead, mice were kept alive until their tumor reached a predefined tumor growth endpoint or for up to 14 weeks after inoculation if the tumor growth endpoint was not reached (**Fig. 2c**). Each mouse was weighed and had their tumor assessed in-person twice per week and underwent a T_2_-weighted MRI once per week. Six of 17 (35.3%) mice across the 300, 600, and LM cohorts (2 mice within each cohort; **Extended Data Table S3**) achieved a complete response (CR), whereas no mice in the NS or NM control cohorts experienced CR (*P* = .0418). For the mice that achieved a CR, the median time to CR from M1 was 14 days (IQR, 14.0-15.5 days), and none demonstrated any evidence of disease until the end of the study at 14 weeks after M0 (example shown in **Fig. 2b**). All control mice (ie, those in the NS and NM cohorts) had consistent tumor growth until meeting the endpoint criteria for euthanasia. All mice within the 300 and 600 cohorts that initially showed a tumor volume decrease or tumor volume stabilization at timepoint M2 but did not achieve a CR (n=8/12) demonstrated evidence of tumor growth during the first weekly T_2_-weighted MRI (ie, 1 week after M2) and then experienced consistent weekly growth until a tumor growth endpoint was met.

There were no observed toxicities or abnormal behaviors noted during the confirmatory study. There was no statistically significant difference in weight changes between cohorts from M0 to M2 (*F*_4,21_ = .063; *P* = .992). All mice gained weight steadily throughout the duration of the study, except for occasional mild-to-moderate weight loss when tumors approached 1000 mm^3^ (**Extended Data Fig. S2**). Importantly, the mice that achieved a CR had a steady and gradual median weight gain of 4.1 g between M0 and week 14. Tissue collection and histological data are presented in **Extended Data Table S5**.

Twenty-two of 27 (300: 5/6, 600: 5/6, LM: 5/5, NS: 4/5, NM:3/5) mice from the confirmatory study had their organs, including the liver, spleen, kidneys, lungs, heart, stomach, and duodenum, collected at the time of euthanasia for histological analysis. Hepatic cytoplasmic vacuolation consistent with glycogen was found in 33.3% (5/15) of mice across the 300, 600, and LM cohorts, but this was not seen in any mice from the NS or NM cohorts. Hepatic cytoplasmic vacuolation consistent with glycogen is an early sign of insulin resistance, which C57BL/6 mice are prone to developing. Of note, the mice in the 300, 600, and LM cohorts had an average survival of 52 days as compared to 27 days for the mice in the NS or NM cohort. All other histological findings within the MENP cohorts were also seen among the control NS cohort with similar frequencies. The six mice that achieved a CR did not show any histological differences in examined tissues compared to their respective cohorts.

### Combined analyses of the pilot and confirmatory in vivo studies

The tumor volume data for the pilot and confirmatory studies are demonstrated in **Fig. 3**, and the relaxometry data are demonstrated in **Fig. 4**. Overall, only the 300 and 600 cohorts experienced consistent anti-tumor therapeutic effects across both studies (**Fig. 3**). Combined tumor volume and relaxivity data are summarized in **Extended Data Table S6.** The ANOVA analyses for the combined data from both studies for absolute T_2_ relaxation changes from baseline yielded *F*_4,38_ = 5.365 (*P* = .002), and relative T_2_ relaxation changes yielded *F*_4,38_ = 7.081 (*P* < .001). The ANOVA analyses for T_2*_ absolute relaxation changes from baseline yielded *F*_4,38_ = 10.708 (*P* < .001), and T_2*_ relative relaxation yielded *F*_4,38_ = 12.873 (*P* < .001). The tumor volume absolute change from baseline yielded *F*_5,21_ = 5.803 (*P* < .001), and tumor volume relative change yielded *F*_5,41_ = 15.271 (*P* < .001).

**Fig. 3:**
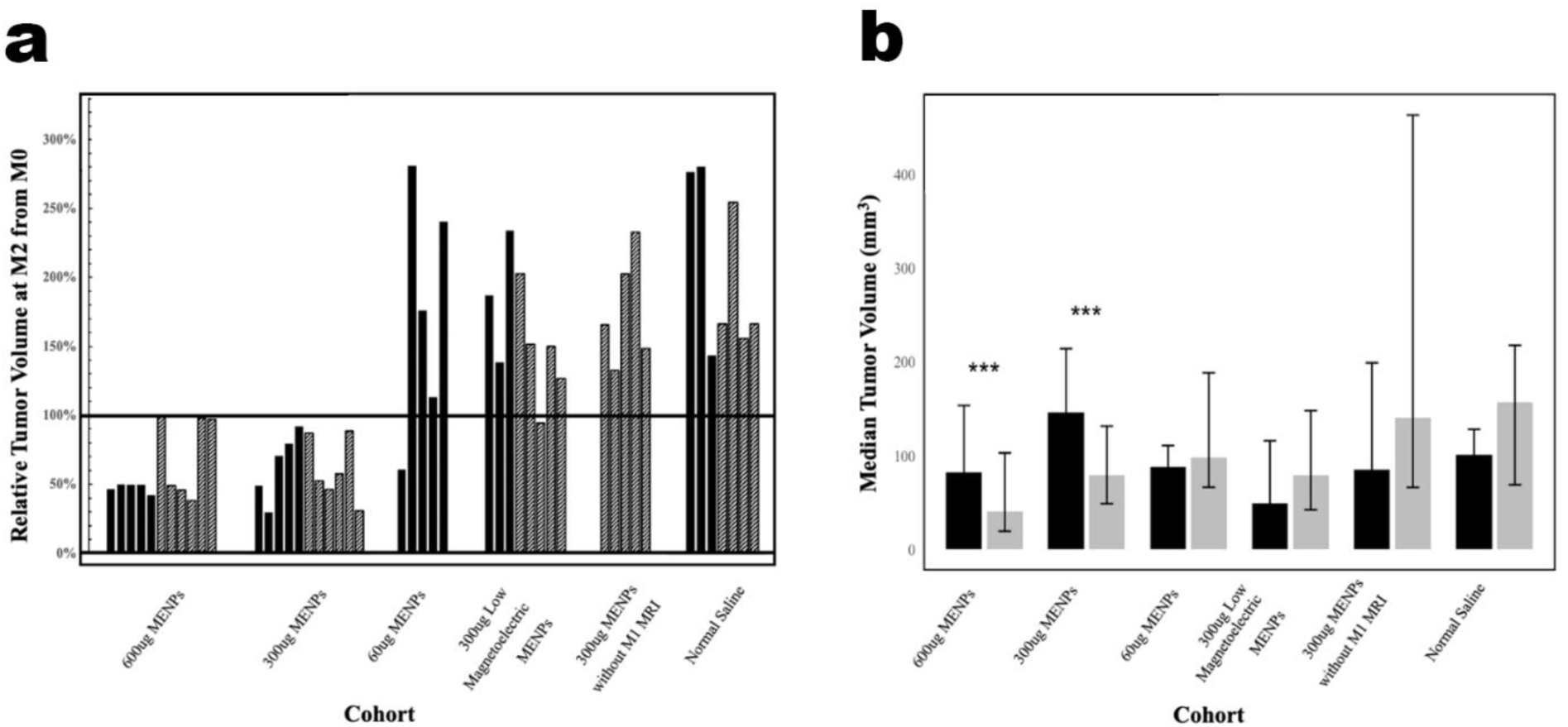
Tumor volume outcomes. **a Clustered bar chart of all mouse tumor volumes at timepoint M2 as a percentage of tumor volume at M0 (ie, at baseline).** Mice are categorized into treatment groups. The mice from the pilot study are represented with solid bars, and mice from the confirmatory study are represented with upward-sloping dashed line bars. No mouse within the 300 or 600 cohorts during either study experienced tumor growth. All mice within the other cohorts experienced tumor growth, except for one mouse in the LM cohort in the confirmatory study and one mouse within the 60 cohort in the pilot study. **b Bar chart of the median tumor absolute volumes for mice in each cohort, both before treatment (M0) and 5 days after treatment (M2).** Both the 600 and 300 cohorts showed a significant decrease in tumor volume following treatment, whereas all other groups show an overall increase in tumor volume. *** represents *P* < .001.

**Fig 4:**
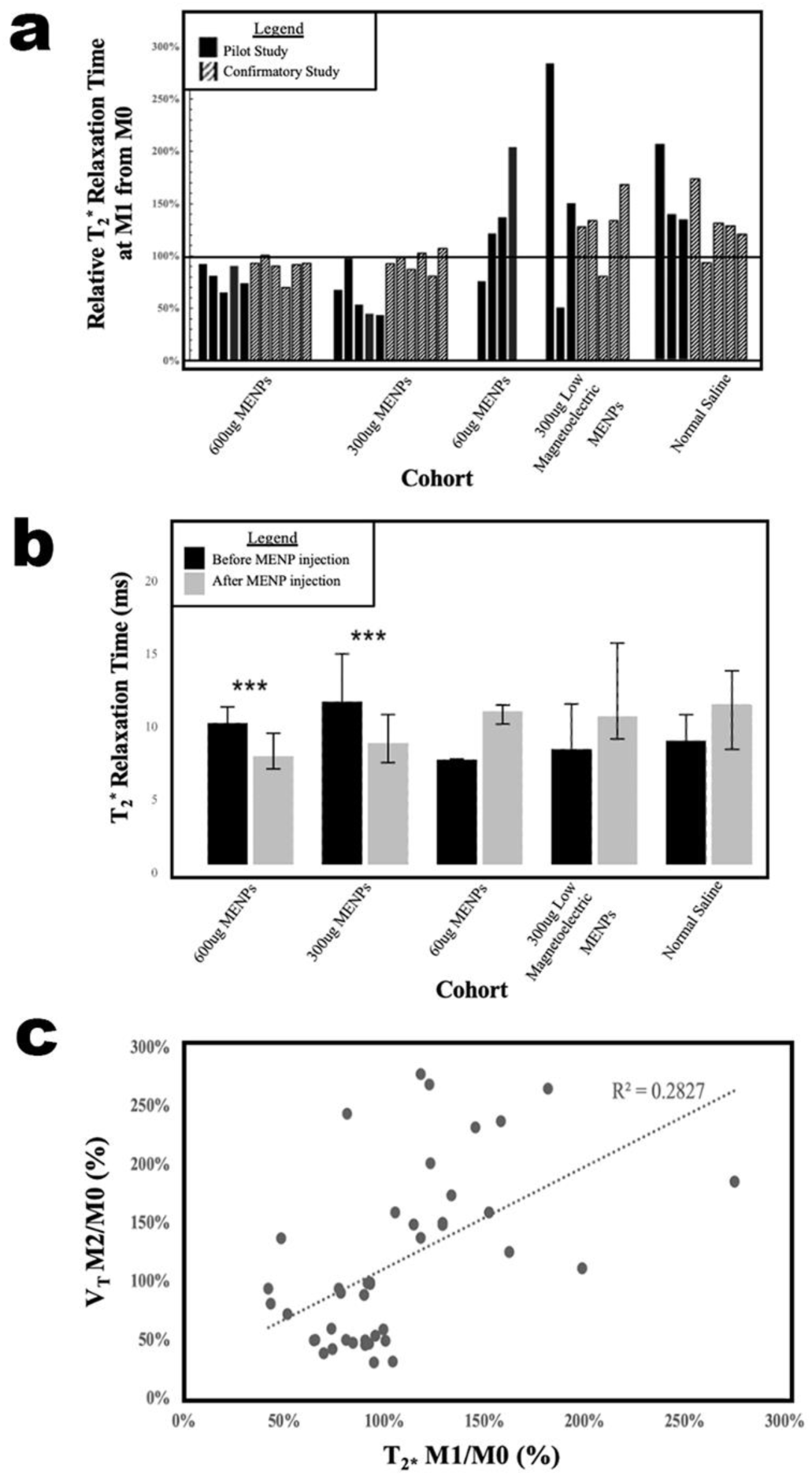
Relaxivity outcomes. **a Clustered bar chart of all mouse tumor T_2_* relaxation times at timepoint M1 as a percentage of tumor T_2_* relaxation times at M0 (ie, at baseline).** Mice are categorized by treatment groups. The mice from the pilot study are represented with solid bars, and mice from the confirmatory study with upward-sloping dashed line bars. **b Bar chart of the median tumor T_2_* relaxation times for mice in each cohort, both before (M0) and after(M1) MENP injection.** ** represents *P*< .01, and * represents *P* < .05. **c Correlation between difference in T_2_* relaxation time and difference in tumor volume across all cohorts**. Scatter plots of relative tumor volume and T_2_* changes at time M2 and M1, respectively, of the pilot and confirmation studies are shown. Tumor volume changes and T_2_* signal modulation are directly correlated (*r*=0.532; *P* < .001 [95% CI, 0.266-0.716]), providing evidence that the degree of T_2_* modulation at time of MRI activation is able to predict tumor response.

LSD analyses for the combined pilot and pilot studies are summarized in **Extended Data Table S7**. The 300 and 600 cohorts demonstrated the greatest relative tumor shrinkage, with reductions of 59.3% and 49.8% from baseline, respectively. In contrast, the NS cohort experienced an increase of 159.4%. Compared to the NS cohort, the mean differences in tumor shrinkage were –133.8% (*P* < .001) for the 300 cohort and –136.7% (*P* < .001) for the 600 cohort. Additionally, the drop in T2* relaxation times at M2 from baseline was 77.4% for the 300 cohort and 67.9% for the 600 cohort, compared to an increase of 121.8% for the NS. Compared to the NS cohort, the mean differences in T_2_* relaxation times were –46% (*P* < .001) for the 300 cohort and –52.3% (*P* < .001) for the 600 cohort. The change in tumor volume was directly correlated with T_2_* relaxometry modulation for both relative (*r*=0.532; *P* < .001 [95% CI, 0.266-0.716]) and absolute (*r*=0.341; *P =* .027 [95% CI, 0.038-0.582]) differences (**Fig. 4**).

### In vitro dye and cell kill studies

Two additional in vitro experiments were performed simultaneously with the pilot and confirmatory studies, with the goals of (i) quantifying the magnitude of the local electric fields generated by the MENPs when exposed to magnetic fields for IRE and (ii) confirming that cancer cells demonstrated evidence of IRE-specific cell-kill (ie, loss of membrane integrity) when exposed to MENPs under the 7 T MRI fields.

The first in vitro study consisted of trypan blue dye degradation experiments to quantify the generation of the MENP local electric fields (**Extended Data Fig. S3**). MENPs were suspended in a trypan blue stock solution and stimulated with an AC magnetic field. The resultant dye degradation was measured using ultraviolet-visible spectrophotometry to determine the ME effect of the particles (**Extended Data Fig. S3c)**. The ME effect was measured with a linear pre-saturation regime. The saturation field ranged between the MENP’s average coercivity field (0.1 T) and the anisotropy field (1 T) depending on the relative orientation of the applied magnetic field and the nanoparticle’s anisotropic magnetocrystalline axes.

MENPs did not have any effect on dye absorption when not stimulated by a magnetic field. Under the presence of a 1 kHz 0.025 T magnetic field, the dye absorption was reduced to 41.9% (*P* = .0002) of the original absorption, corresponding to an ME effect of 1 VA^26^. LMs also did not show a significant effect on dye absorption without stimulation but reduced absorption to 64.8% (*P* = .0115) after magnetic stimulation, corresponding to an ME coefficient of 0.5 VA (equivalent to 150 µg of MENPs with a ME coefficient of 1 VA; **Extended Data Fig. S3c**). The membranes of SKOV-3 cells treated with MENPs and simultaneous magnetic field stimulation were permeabilized, as demonstrated by propidium iodide fluorescence under microscopy (**Extended Data Fig. S3d**).

The second in vitro study consisted of 6 samples of KPC961, 3 of which were treated with MENPs and 3 control samples that did not receive MENPs but were exposed to the same 7 T MRI relaxometry fields as in the in vivo studies. One hour after MRI exposure, all samples underwent staining with annexin V and propidium iodide (PI) followed by flow cytometry. Early apoptosis, late apoptosis, and necrotic cell death were then compared between the experimental and control samples. The MENP samples demonstrated a statistically significant increase in both early apoptosis (4.5% vs 2.4%; *P* < .022) and late apoptosis (7% vs 5.3%; *P* < .006; **Fig. 5a**). To characterize MRI-activated MENP cellular death as a function of time, this procedure was repeated twice for the 3 MENP samples at 2 and 3 hours after MRI exposure. The data demonstrated increases in both early and late apoptosis as a function of time after MENP MRI exposure (**Fig. 5b**). An HMGB1 assay was used to determine if a there was relationship between HMGB1 concentration and a combination of particle dose and field strength (**Fig. 5c**), as HMGB1 has previously been demonstrated to be liberated during electroporation and electrochemotherapy^33^. Additionally, trypan blue was used to measure the reduction in cell viability and indicated a 20% reduction in viability after treatment (**Fig. 5d).**

**Fig. 5:**
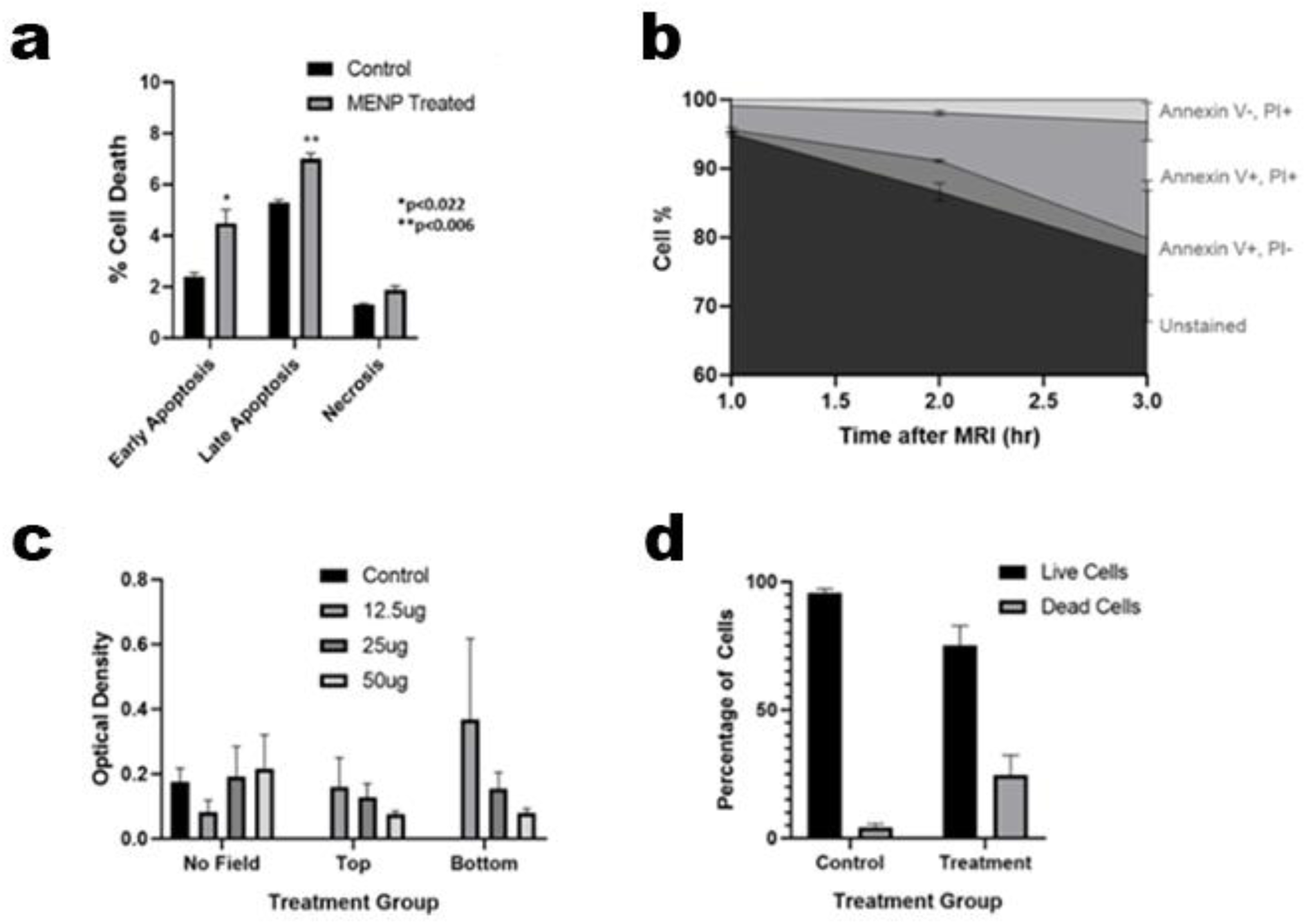
In vitro cell death mechanism data from KPC961 cells treated with MENPs and magnetic stimulation. **a Percentage of KPC961 cells killed through early apoptosis, late apoptosis, and necrosis.** Cell kill was measured via flow cytometry through annexin V and PI staining. All cells underwent the same studies as the in vivo experiments on a 7 T MRI and were then stained with 5 µl of annexin V and 10 µl of PI to test for late and early apoptosis, respectively. The MENP samples demonstrated a statistically significant increase in both early apoptosis (4.5% vs 2.4% for controls; *P* < .022) and late apoptosis (7.0% vs 5.3%; *P* < .006), but no significant change in necrosis rates were seen. **b Annexin V and PI uptake over time after treatment with MENPs and MRI.** A steady decrease in unstained cells demonstrated cell death over 3 hours after treatment. An increase in Annexin V–positive and PI-positive cells over time is indicative of apoptotic cells with permeable membranes. **c HMGB1 assay**. Results are shown for KPC961 cells treated with various doses of MENPs and DC magnetic stimulation using a permanent magnetic placed either directly under the cell dish (bottom) or with a 1.5cm gap (top). Two-way ANOVA suggests a significant interaction between particle dose and field strength (*P* = .0292) on HMGB1 concentration. **d Cell death percentage as measured by Trypan Blue assay**. MENP samples demonstrated a significant increase in dead cells (*P* = .0097).

## Discussion

This study presents the first in vivo and in vitro evidence of an externally controlled, targeted, predictive, MRI-based, theranostic agent capable of inducing IRE within a solid tumor. A pilot study of 21 mice and a confirmatory study of 27 mice demonstrated that ≥300 µg doses of MENPs achieved a mean three-fold reduction in tumor volume (62.3% vs 188.7%; *P* < .001) over 5 days from MENP activation (ie, when exposed to MRI magnetic fields). MENPs also demonstrated predictive theranostic potential, with the drop in T_2_ relaxation time on MRI directly correlating to tumor volume reduction (*r*=0.351; *P* = .039). Additionally, 6 of the 17 (35.3%) mice in the experimental MENP arms in the confirmation study achieved a durable CR.

MENPs achieved an approximate three-fold reduction in tumor volume when used in doses of 300 µg (59.3% reduction) and 600 µg (49.8%) compared with NS controls (159.4% tumor volume increase; *P* < .001) when exposed to MRI magnetic fields. Furthermore, no mice that received a dose of 300 or 600 µg experienced tumor growth at 5 days after MENP exposure to MRI magnetic fields. MENPs also demonstrated strong signal modulation on T_2_ and T_2*_ MRI relaxometry, with the relaxation times of 77.4% for the 300 cohort and 67.9% for the 600 cohort compared to 121.8% for the NS cohort (*P* < .001). Additionally, T_2*_ signal modulation directly correlated to tumor volume reduction (*r*=0.532; *P* < .001 [95% CI, 0.266-0.716]), meaning that it appears possible to predict tumor response as a function of signal modulation at the time of treatment. In the confirmatory study’s MENP cohorts (ie, the 300, 600, and LM cohorts), 35.3% of mice achieved a durable CR with a single MRI-activated MENP dose as compared to NS and NM controls (*P* = .0418). In contrast, all mice from the NS and NM control cohorts experienced continuous tumor growth until they met euthanasia endpoints. Lastly, there were no discernable MENP-related toxicities or substantial weight loss reported for any mice at any timepoint. On histological analysis, the only change seen within mice from the experimental arms of the confirmatory study was hepatic cytoplasmic vacuolation consistent with glycogen.

The ME effect distinguishes MENPs from all other nanoparticles and allows efficient coupling of magnetic and electric fields, thus unlocking the theranostic capabilities described herein. Unlike magnetic fields, electric fields can have dramatic effects on fundamental biological processes, as is seen with the conventional IRE process; however, electric fields cannot be wirelessly/noninvasively transferred to or from any desired internal target site within the biological microenvironment, thus requiring the use of wired electrodes. In contrast, unlike electric fields, magnetic fields can propagate through both conductive and dielectric biological tissues without dissipation or scattering, eliminating the need for physical electrodes to induce strong local electric fields at targeted sites.

Furthermore, MENPs exhibit preferential targeting for cancer cells due to the increased conductivity and non-polarity of cancer cellular membranes, which distinguish them from normal tissue^34^. These characteristics of cancer cells are directly produced by the dysregulation of cell cycles and metabolism, which are key features of cancer across cancer types^35^. As such, the specificity of this preferential targeting mechanism may be agnostic to solid tumor histology^16,27^. Through the combination of this unique cancer cell–specific targeting mechanism and the generation of highly localized electric fields upon application of an external magnetic field, MENPs can induce a unique form of highly ablative IRE while simultaneously minimizing any harmful effects to surrounding normal tissues.

Cancer cells are vulnerable to this MENP targeting mechanism because their rapid proliferation creates membranes with relatively high electrical conductivity^27^. According to current consensus, this high electric conductivity might lower their membrane electroporation threshold by 20 to 100 times that of normal tissue^36^. Because of their nanoscale size and the ME effect, MENPs leverage this difference to specifically target cancer cells. Given that the nanoparticles’ electric fields are activated upon application of a relatively small magnetic field (eg, a 1.5 kOe field generated by placing a permanent neodymium magnet over a flank tumor), the net force guides MENPs towards the highly conductive cancer cells and away from the less conductive surrounding normal cells^16,27^. Once the MENPs have been targeted to the tumor, application of a relatively strong MRI magnetic field causes MENPs to generate a local electric field strong enough to electroporate the cell membrane in the immediate vicinity of the nanoparticles^15,27^.

In this study, we presented the first evidence of MENP-based IRE. Although previous studies demonstrated that MENPs can at least induce reversible electroporation during targeting^37^, a clear and reliable ablative IRE effect has not yet been shown. Our in vitro dye experiment demonstrated that MENPs have a magnetic-to-electric field conversion on the order of 1 VA. Given the saturation limit of the MENPs identified via the linear Landau theory^38^, MENPs can generate local electric fields of over 1000 kV/m across the cellular membrane if they are in direct contact with the cell^30^ while activated by an MRI magnetic field of ≥1 T. This occurs because a magnetic field of this strength (ie, ≥1 T) can exceed the MENPs’ magnetocrystalline anisotropy field, H_K,_ thus achieving the strongest possible local electric field that a MENP is capable of generating. This explains why the neodymium magnet, which is used for the targeting of the MENPs and only generates a magnetic field of 0.15 T, was not strong enough to trigger an MENP-based IRE effect (**Extended Data Fig. S3b**). However, MRIs with a field strength of ≥1 T are capable of triggering an MENP-based IRE^39^. When applied to MENPs within the vicinity of cancer cells, the MRI field increases both early apoptosis (4.5% vs 2.4% for controls; *P* < 0.022) and late apoptosis (7.0% vs 5.3%; *P* < .006), both of which increase as a function of time from MRI (**Fig. 5b**). Additionally, the associated increase of HMGB1 (**Fig. 5c**) and reduction in cell viability (**Fig. 5d**) also suggests that IRE plays an important role in the particles’ ablative cell kill upon activation. However, it is important to note that there are likely multiple interrelated mechanisms that ultimately cause tumor cell kill upon MENP activation. We used dye degradation as an indirect measure of a local electric field strength via the generation of oxidative free radicals that react and degrade the organic dyes (**Fig S3c**). Therefore, it is likely free radical generation during MENP activation likely contributes to in vivo cell kill, as it does with conventional IRE^40^.

The mechanisms of how MENP-based IRE cause cell death are unlikely to precisely mirror those of conventional IRE techniques. Conventional IRE uses kilovolts of voltage and frequencies up to hundreds of kilohertz to overcome the screening effects of the conductive intracellular/extracellular space and deliver sufficient energy to the cell membranes. In contrast, MENP treatment applies the electric field directly to the membrane surface and avoids the destructive screening effects within the relatively highly conductive intracellular and extracellular spaces, so the required amount of energy to achieve the threshold (breakdown) transmembrane potential is much smaller^41^ (**Extended Data Fig. S6**). As such, while conventional IRE offers a useful guide for expected results and mechanisms of actions, the full breath of cellular effects that MENP-based ablation induce is likely to differ in important but currently unknown ways. MENP-based IRE is highly localized within the membrane, with a lateral size defined by the nanoparticle size (∼30 nm); thus, it is substantially more efficient and has significantly fewer side effects than the conventional IRE probe-based approach^23–25^.

MENPs can also be used as T_2_ contrast agents by shortening the spin-spin time constant^42^, which generates a quantifiable negative contrast in the image^43^. MENPs induce MRI signal modulation by decreasing spin-spin (ie, T_2_) relaxation times because of their nonzero-magnetic-moment cores. However, T_2_* is a more sensitive mapping study mechanism for detecting MENPs in vivo because of their ferrimagnetic cores^44^. MENPs have been previously characterized with in vitro and ex vivo phantom studies to determine T_2_* signal modulation as a function of concentration^32^; signal intensity deltas increased with higher concentrations of MENPs in a linear relationship with doses up to 500 µg, which was the highest tested dose^32^. In a healthy baboon model, MENPs were detectable due to a reduction in T_2_* relaxation times within organs that they preferentially accumulated within, such as the liver, kidneys, and spleen^32,45^.

For the first time, we demonstrated that MENPs are intratumorally detectable via MRI. This effect was best seen at the 300 µg and 600 µg doses, which demonstrated a reduction of T_2*_ relaxation by 2.2 and 2.5 ms, respectively. Although this effect is modest compared with other common MRI contrast agents, this feature could be potentially paradigm-shifting when considered in combination with MENPs’ therapeutic predictive power. Notably, the degree of intratumoral signal modulation was directly correlated to the degree of tumor response (*r* = .656; *P* < .001 [95% CI, 0.455-0.794]). This increased signal modulation likely reflects a greater degree of targeted MENP concentration within the tumors, as has been previously demonstrated in phantom studies^32^, which ultimately translates into more complete cell-kill due to MRI-activated MENP-IRE. However, the inherent complexity of imaging a tumor while it is actively being exposed to an ablative therapy poses several limitations. One of the major issues is that our mouse model is immunocompetent, meaning that there was likely an acute inflammatory response occurring during the time of imaging, which could increase T_2_/T * relaxation time due to increased edema within the TME. Because an MRI’s magnetic field is always active, the particles would theoretically be activated as soon as the mouse was inserted into the machine during set up; this competing inflammatory reaction, due to rapid initial IRE cell kill and generation of reactive oxygen species (ROS), likely masks the strength of the relaxivity modulation seen with the MENPs. This phenomenon may help explain why the 60 cohort demonstrated an overall increase in T_2_/T_2_* relaxation time from baseline as compared to the 300 and 600 cohorts (**Fig 4b**). Beyond an acute inflammatory reaction, it is also unclear how rapid cell-kill within the TME would ultimately contribute to relaxivity modulation in real-time.

Despite these challenges, MENPs still demonstrated an overall significant effect on relaxivity that directly correlated with tumor volumes 5 days after activation. This relation underlies MENPs’ ability to operate as a first-of-its-kind anti-cancer theranostic nanotherapeutic agent. The real-time predictive adaptive therapy enabled by MENPs would allow for oncologists to adjust treatment plans in real time; thus, MENPs may play an important role in the rapidly developing field of personalized cancer care by enabling real-time tumor- and patient-specific treatment approaches. Moving forward, radiomic modeling and machine learning tools may prove to be more powerful methods to fully exploit MENP theragnosis.

Although inorganic NPs have been reported to cause acute and late inflammatory toxicities^46^, no dose-limiting toxicities have been reported with any of the currently published MENP in vivo studies^16,32^. MENPs have been found to be nontoxic at relatively high concentrations and are excreted from murine models in approximately 2 months^32,47^, with >90% clearance in 2 weeks^16^. Doses as high as 500 µg in a non-human primate model have demonstrated no signs of toxicity^32^. In line with these results, we did not observe any discernable toxicities among mice in the 300 or 600 µg cohorts. Researchers have previously demonstrated through Raman histopathology that MENPs do not appear to damage any normal bodily tissues (eg, liver, lung, bladder, spleen, kidneys, adrenal gland, stomach, uterus, bowel, heart, or CNS tissue) and confirm effective hepatobiliary and renal excretion within ∼30 days^32^. MENPs also did not seem to affect blood chemistry within a 30-day interval from administration^32^. Our study is congruent with these prior studies and highlights the apparent favorable safety of MENPs, as there were no discernable toxicities associated with MENPs at any dose level. Although there was evidence of tissue abnormalities upon histologic examination (**Extended Data Table S7**), these toxicities were also seen at similar rates within the NS control cohort. Hepatic cytoplasmic vacuolation was the only histological finding that was only seen within the MENP experimental arms (33.3% of MENP-treated mice vs 0% of control mice). However, C57BL/6 mice are known to be particularly susceptible to developing insulin resistance, and this finding likely represents early insulin resistance changes as a result of the longer average survival times for mice in the experimental vs control arms (52 days vs 27 days, respectively).

Though this study provides the first in vivo evidence of an external field–controlled, predictive, MRI-based theranostic agent for targeting and treating solid tumors through IRE, it has several limitations. First, the flank model limits the generalizability of these results, as most tumors cannot be targeted in such a simple manner. Second, the study only focused on the tumor response after a single dose; it is unclear whether multiple doses of MENPs may have additive effects or whether additional doses will continue to be predictive of tumor volumetric response. Third, we used an MRI unit with 7 T, and it is unclear whether the relaxometry modulation effects would be detectable under the more common clinical MRI field strengths of 1.5 to 3 T (**Extended Data Fig. S3b**). Fourth and finally, we did not investigate how the additional magnetic fields within the MRI (ie, the gradient and radiofrequency fields) influence the MENPs’ interactions with cellular membranes, and it is unclear whether these fields are necessary for MENPs to realize their full therapeutic potential. These limitations underscore the need for further preclinical and clinical studies to better understand and refine the use of MENPs as a unique theranostic tool within the oncological landscape.

## Conclusion

This study is the first to demonstrate an innovative application of MENPs as an external, magnetic field–controlled, MRI-based, theranostic agent capable of targeting and ablating tumors via IRE in an in vivo solid tumor model. A significant proportion of mice experienced a durable CR from a single dose of MENPs. MENPs can potentially serve as a predictive theranostic agents because their signal modulatory effect is directly correlated to their anti-tumor therapeutic effect. These capabilities underscore the unique potential of MENPs as a new modality for real-time adaptive therapy. Future studies must explore multi-dose approaches, orthotopic models, and immunomodulation to better understand the role MENPs may have in human oncologic care.

## Materials and Methods

### MENP synthesis

MENP synthesis was based on a recently described process that demonstrated 1 VA of the ME effect using a nanoprobe-scanning tunneling spectroscopy technique at a single-nanoparticle level^26^. To further optimize the MENPs for this study, in conjunction with the aforementioned nanoprobe measurement as a calibration technique, we used a special dye degradation test to evaluate the strength of the ME effect of all the nanoparticle types used in vitro and in vivo as well as the cores of the nanoparticles (**Extended Data Fig. S1C).** MENPs were synthesized from the following chemicals: cobalt(II) nitrate hexahydrate (Co(NO3)2•6H2O) (Sigma Aldrich, St Louis, MO), iron(III) nitrate nonahydrate (Fe(NO3)2•9H2O) (Sigma Aldrich), sodium hydroxide (NaOH; Sigma Aldrich), barium carbonate (BaCO3) (ThermoFisher Scientific, Waltham, MA), titanium(IV) isoproproxide (Ti[OCH(CH3)2]4) (Sigma Aldrich), citric acid (HOC(COOH)(CH2COOH)2) (Sigma Aldrich), and ethanol (>99.7%) (Millipore Sigma, Burlington, MA). All reagents were used without further purification.

Core-shell MENPs were synthesized using a modified designed to promote anisotropic growth^11,26,48^. The cobalt ferrite cores were fabricated via a coprecipitation process using metal salts of cobalt and iron, with sodium hydroxide as the precipitating agent. In a typical synthesis, 100 mg of cobalt nitrate and 278 mg of iron nitrate were dissolved with constant stirring into separate beakers containing 20 mL of deionized (DI) water. The beakers were brought close to the reaction temperature of 90°C and then mixed together. A 3M sodium hydroxide aqueous solution was added until the mixture reached a pH of 13, at which point the precipitation process started. The reaction was allowed to evolve for 1 hour to reach 25 to 30 nm particle size. The solution was then cooled and the cobalt ferrite nanoparticles were magnetically separated from the solution. The nanoparticles were then washed twice in DI water and once in ethanol to remove excess reactants before drying overnight on an 80 °C hotplate.

The barium titanate shells were formed on the cobalt ferrite cores at a 1:2 (core:shell) stochiometric ratio using a modified solgel and auto-combustion process. For synthesis, 50 mg of the previously prepared cobalt ferrite cores were mixed in 20 mL of DI water with 1000 mg of citric acid. This beaker was then probe sonicated for 2 hours to fully disperse the cores. In separate beakers, 88 mg of barium carbonate was mixed with 20 mL of DI water, and 126 μL of titanium isopropoxide was mixed with 20 ml of ethanol. Next, 1000 mg of citric acid was added to both the barium carbonate and the titanium isopropoxide beakers and allowed to mix for 1 hour at room temperature to ensure full chelation. These beakers were then combined, and the resultant solution was brought to 90 °C to begin evaporating the excess ethanol and DI water. When the combined barium and titanium solution reached 20 mL, the sonicated cobalt ferrite cores were added. The final solution was then allowed to evaporate until a gel formed. The gel was then slowly heated to and held at 800 °C and for 6 hours before slowly cooling to room temperature. The final product was ground down in a mortar and pestle. This was then washed twice in DI water and once in ethanol before drying overnight at 80 °C.

Low-ME-effect control nanoparticles (LM) were synthesized similarly to the core-shell MENPs. However, during the core addition step, the dispersion step was kept shorter (0.5×) to promote multi-core agglomerates, thus effectively reducing the number of single cores. These multicore agglomerates have poorer lattice matching between the core and shell due to misaligned grain directions, and the resultant ME effect, according to the described dye test, is effectively reduced by a factor of 2.

### Nanoparticle characterization

The magnetic properties of the nanoparticles were measured using a MicroMag 2900 Alternating Gradient Magnetometer (Lakeshore, Carson, CA). M-H loops were made from either ±0.5 or ±1 T sweeps to achieve saturation. The size of the nanoparticles was verified using a multimode atomic force microscope (Bruker), a magnetic force microscope (Bruker), a high-resolution transmission electron microscope (JEOL JEM-2100 Plus with a 200 kV LaB6 electron source; Jeol USA, Peabody, MA), and dynamic light scattering measurements. The elemental composition was independently confirmed through energy-dispersive spectroscopy and x-ray diffraction analyses^26,49^.

### Dye degradation tests

To further optimize and test the efficacy of the MENPs, their ability to degrade dye was used as a proxy to measure the strength of their ME effect. We used earlier comprehensive measurements of the ME effect at a single-nanoparticle level with the nanoprobe scanning tunneling spectroscopy mode of a scanning probe microscope (Bruker) as a calibration value^26^. First, a diluted Trypan blue solution was prepared by mixing the commercial 0.4% Trypan Blue Solution (ThermoFischer Scientific) with DI water to create a 0.04% Trypan blue stock solution. Five mg of MENPs were weighed out and added to 1 mL of the Trypan blue stock solution. The mixture was then sonicated for 1 hour to disperse the MENPs and reach an adsorption/desorption equilibrium of the dye-MENP mixture.

The dye samples were then divided into 2 groups: one for sonication treatment and one for magnetic stimulation. The sonication group samples were probe sonicated for an additional hour. The magnetic stimulation groups were also probe sonicated for an hour, but during this time, they were exposed to a 1 kHz, 0.025 T alternating magnetic field.

After the respective treatments, the samples were centrifuged to remove the MENPs. The supernatants were further diluted with DI water in a 1:4 ratio, and their absorbance spectra were measured using an ultraviolet-visible spectrophotometer (Shimadzu UV-Visible Spectrophotometer UV-2600; Shimadzu Scientific Instruments, Columbia, MD). The peak absorbance at around 580 nm for each treatment group was compared to an untreated Trypan blue stock solution control sample. Each test was performed in triplicate.

### Nanoparticle solution preparations

We have previously shown that it is critical that MENPs are in direct contact with the cellular membrane surface in order to leverage the efficient wireless induction of a local electric field^30^. As soon as the nanoparticles are removed from the dielectric membrane surface, the electric field drops from over 100 to below 0.1 kV/m because the MENPs’ electric fields are screened out by free ions in the conductive extracellular and intracellular spaces, with the Debye length in the sub– 1-nm range^50^. Therefore, to ensure that the nanoparticles preferentially target the membrane, MENPs were coated with a thin layer of polyethylene glycol (PEG)^11^.

The PEG coating conditions were chosen to control the balance between attraction forces (such as the van der Waals interaction and the MENPs’ dipole–cellular membrane force) and repelling forces (such as the Coulomb force, which is caused by surface charge distributions on the membrane and nanoparticle surfaces) between the nanoparticles and the cellular membrane^27,50^. PEGylation also improved MENP solubility and decreased immunogenicity. The nanoparticles were coated in PEG (H(OCH2CH2)nOH) (Sigma Aldrich) with a molecular weight of 600Da. MENPs were weighed out and transferred to a 15 mL tube. DI water was added at a 1:1 ratio, followed by sonication in a Q125 Probe Sonicator (QSonica, Newton Borough, CT) for 2 hours using a 3-second on/3-second off cycle at an amplitude of 20%. Then, 600Da of liquid PEG was introduced at approximately 8% of the total volume and sonicated for an additional 2 hours. The solution was then washed 3 times with DI water using centrifugation at 3000 relative centrifugal force (rcf) for 7 minutes. The nanoparticles were redistributed in DI water such that the ratio of MENPs to solution was 2:1. Three hundred μL of this solution per mouse were set aside for the 600 μg cohort. Thereafter, DI water was added to the remaining solution to double the volume and obtain a 1:1 ratio of MENPs to DI water. Three hundred μL of this solution per mouse was set aside for the 300 μg cohort. LM were PEGylated separately in the same manner and were redistributed in DI water such that the ratio of MENPs to solution was 1:1. Then, 300 μL of this solution was set aside per mouse in the LM cohort. The MENPs were then magnetized using a permanent magnet, generating a 0.15 T field, and sonicated in their individual vials until the time of injection (minimum of 30 minutes) for optimal dispersion.

### Outline of in vivo murine flank tumor experiments

Two murine experiments using a flank tumor model were performed in succession to test the theranostic potential of MENPs and evaluate the in vivo effects of MENPs in conjunction with MRIs regarding tumor-cell kill and MRI relaxometry. Both studies followed the same protocol (**Fig. 2**) regarding particle-solution preparation and timing of baseline MRIs (timepoint M0), MRIs to evaluate for T_2_ contrast effects and trigger tumor cell kill (timepoint M1; delivered 48 hours after M0 and approximately 16 hours after tail-vein injections), and MRIs to evaluate for tumor cell kill (timepoint M2; 5 days after M1/7 days after M0). The primary aims for the pilot study were (i) differences in signal modulation of T_2/_T_2_* MRI mapping studies between M1 and M0 and (ii) differences in MRI tumor volume between M2 and M0. For the pilot experiment, mice were immediately euthanized after their M2 MRI.

The follow-up confirmatory study cohort design was informed by the pilot experiment. The primary aims for the confirmation study were the same as the pilot study, with the addition of characterizing tumor volume response over time and histological analyses of organs to evaluate the effect of the particles within vital bodily organs. Normal tissue collection was performed at the time of euthanasia. For the confirmation study, the study endpoint was 14 weeks after M1, at which point the mice were euthanized. To minimize morbidity, mice were euthanized before 14 weeks if they experienced (i) tumor volume ≥1,000 mm^3^, (ii) tumor diameter ≥1 cm, or (iii) tumor ulcer diameter ≥5 mm.

### Murine pancreatic ductal adenocarcinoma culture and murine model

In vitro and in vivo experiments were conducted using the UN-KPC-961 pancreatic cell line. This cell line was obtained via a material transfer agreement from Dr. Surinder K Batra (University of Nebraska Medical Center, Omaha, NE). Cells were maintained and cultured according to their suggested protocols using DMEM:F12 supplemented with 10% heat-inactivated fetal bovine serum (FBS) and penicillin-streptomycin (100 units/mL–100 mg/mL). All cells were maintained under standard tissue culture conditions at 37 °C and 5% CO_2_ and were tested and kept mycoplasma-free. All steps were performed under sterile conditions in a biosafety cabinet.

All mice were maintained under Institutional Animal Care and Use Committee guidelines. Eight-to ten-week–old C57BL/6 mice (The Jackson Laboratory, Bar Harbor, ME, USA) were used as hosts for the KPC-961 cells. One million KPC-961 cells were subcutaneously injected into the flanks of C57BL/6 mice at a volume of 100 µL/mouse in 1× phosphate-buffered saline (PBS).

### In vitro and in vivo MRI scans

All MRI scans were conducted with a horizontal 7 T Biospec MRI (Bruker) using a 35 mm birdcage coil (Doty Scientific, Richland County, SC). Mice were anesthetized with 2% isoflurane in O_2_ and restrained in a specific holder during data acquisition. Anatomical T_2_-weighted axial images of the abdominal cavity were obtained with a turbo spin echo rapid acquisition with relaxation enhancement (TurboRARE) sequence with repetition time (T_R_)/time to echo (T_E_) of 4000/48 ms, 30 slices, and 1 mm of slice thickness (no gap between slices). The field of view (FOV) was 32×32 mm and 256×256 pixels. T_1_, T_2_, and T_2*_ maps were acquired using the same slice details, FOV, and image resolution with a multi-echo TurboRARE sequence (T_R_ = 5000 ms, 16 echoes with 7.52 ms echo spacing to increase T_E_ from 7.52 ms to 160.32 ms). Mapping studies were always performed in the following order: 1) T_2*_ mapping, 2) T_2_ mapping, and 3) T_1_ mapping. All scans were performed during the same session. The mouse was exposed to the static magnetic field for approximately 40 to 50 minutes per mapping session and for 15 to 20 minutes per TurboRARE imaging session.

### In vitro apoptosis assay by flow cytometry

We used the apoptosis cell death assay (Annexin V/PI) kit (cat#640914; BioLegend, San Diego, CA). One million mouse pancreatic cancer cells from the Kras^G12D^/Trp53^R172H^/Pdx-1-Cre (KPC) cell line were combined with 100 µl of cold Annexin V binding buffer. This mixture was then transferred to 5 ml flow cytometry tubes with either control cells (n = 3 tubes) or MNEP-treated cells (n = 3 tubes). Then, 5 µl of Annexin V FITC and 10 µl of PI were added to each tube. Tubes were gently vortexed and incubated at room temperature for 15 minutes in the dark. After incubation, 400 µl of Annexin V binding buffer was added to each tube. The stained samples were then analyzed on a FACSort flow cytometer (BD Biosciences, Franklin Lakes, NJ). The data were analyzed using the FlowJo software (BD Biosciences).

### Tumor segmentation on MRI

Tumor segmentation was performed on coronal slices for every MRI scan via 3D Slicer^51^ v.5.4.0 (open source). The T_2_ relaxation time map was used as the primary series for tumor volume segmentation during the mapping studies, although other series were used as needed to clarify tumor extent if there was significant signal dropout at the periphery. The segmentation statistics tool was used to derive volumetric and average relaxation time data for each contour. All MRI segments used to determine tumor volume underwent blind review by a board-certified diagnostic radiologist. Imaging artifacts (eg, breathing motions, volume averaging, and susceptibility artifacts) can have significant impact on average relaxation time and are primarily located on the periphery of the volume segment; to control for this, an additional segment was derived from the volume segment by performing a 0.8 mm to 1.2 mm contraction (**Extended Data Fig. S4**). If any artifact remained within a small portion of the contracted segment, an additional manual edit was performed to eliminate it. On some M1 contours, removal of the magnet caused mild epilation that appeared as an edematous bright signal, which was also intentionally eliminated from the contour via 0.8 mm to 1.2 mm contraction. If needed, an additional manual edit was performed to ensure that the segmented sample was the best representation of the tumor parenchyma.

### In vivo tumor volume and relaxation time endpoint definitions

Relaxation time endpoints (reflecting the contrast effect) were calculated based on the ratio (presented as percentages) of M1 to M0, and tumor volume endpoints (indicating the therapeutic effect) were calculated based on the ratios of M2 to M0. A CR was defined as an absence of tumor on imaging and at the bi-weekly physical examination. The date of CR was recorded as the time of MRI when the tumor was first noted as absent. All mice with a CR still underwent bi-weekly physical examination and weekly MRIs.

### Definitions for mouse euthanasia endpoints

Tumor endpoints that triggered euthanasia were tumor volume ≥1,000 mm^3^, tumor diameter ≥1 cm, or tumor ulcer diameter ≥5 mm. Tumor volume measurements were performed using MRIs (performed at least once per week), tumor diameter measurements were performed either via MRI or calipers, and tumor ulceration measurements were performed with calipers. Additional protocol endpoints for euthanasia were >20% weight loss from baseline, decreased mobility due to tumor morbidity, and rapidly decreasing performance status; however, none of these endpoints were met by any mouse.

### Tissue preparation for histological analysis

Mice were euthanized by means of CO_2_ inhalation. Immediately after euthanasia, the tissues of interest were excised and stored in a 10% formalin solution overnight at room temperature. Tissues were processed using the VIP E 300 Tissue Tek Tissue processor SN 48940652 (Sakura Finetek USA, Torrance, CA) to create fixed, dehydrated, and cleaned cassettes; femur bone marrow samples were decalcified prior to slide preparation. Tissues were transferred to warm paraffin-filled molds using warm forceps and allowed to cool until solid. A Leica 2125 Microtome (Leica Biosystems, Wetzlar, Germany) was used to cut 4-μm sections, which were then flattened by floating in a 40 °C water bath before being placed on VWR Superfrost Plus microscope slides (VWR, Radnor, PA) to dry. Coverslips were added using Tissue Tek SCA Automated Coverslipper (Sakura Finetek USA) either at this point or after hematoxylin and eosin (H&E) staining using standard procedures (ie, using the MMI H&E Staining Kit Plus [Molecular Machines & Industries, Eching, Germany]). The sectioned tissue slides were imaged using Olympus BX51 Fluorescence Microscope (Olympus Life Science, Tokyo, Japan) at 20× and 40× magnification.

### Histological analyses and pathology quality control

Analyses of selected tissue types (bone marrow [from femur], stomach, duodenum, heart, kidneys, liver, and lungs) were performed on H&E-stained tissue sections . Histological tissue damage evaluation for toxicity was performed by the Tissue Core at Moffitt Cancer Center, which is College of American Pathology– (CAP-) accredited. Histopathological evaluations were performed in accordance with the guidelines of the National Toxicology Program from the US Department of Health and Human Services.

A comprehensive scoring system was created by combining the National Toxicology Program’s Nonneoplastic Lesion Atlas^52^ as a scientific approach with project-centered research model analyses. Histologic findings from H&E-stained slides of each tissue type (bone marrow, stomach, duodenum, heart, kidneys, liver and lungs) were scored as 1 (mild), 2 (moderate), or 3 (severe) based on the most common cellular damage on each tissue type/organ.

### Statistical analyses

Statistical tests were conducted using SPSS version 29 (IBM, Armonk, NY). Planned ANOVA tests were used to evaluate baseline cohort differences after randomization for tumor volume and weight data. Planned ANOVA and LSD tests were used to evaluate relative and absolute tumor volume and relaxation time outcome data from baseline and compare between cohorts for the pilot study, confirmatory study, and combined analysis. For the pilot study, planned student *t* tests were used to determine tumor volume differences between sexes. Planned Pearson product-moment correlation tests were used to evaluate changes in tumor volume and changes in relaxation time for the combined analysis. Planned statistical significance was defined as a two-sided *P* < .05 for ANOVA, LSD, and Pearson product-moment correlation tests. A Fisher exact test was used to compare CR data between the experimental and control cohorts, with statistical significance defined as a one-sided *P* < .05. Ad-hoc analyses comparing differences of grouped cohorts (ie, experimental vs control arms) were performed via student *t* tests, with statistical significance defined as a two-sided *P* < .05. For the dye degradation tests, ANOVA analyses were used to determine whether there were statistically significant differences between groups, and Kruskal-Wallis tests were used to compare groups with a significance level of 0.05.

## Acknowledgements

Editorial assistance was provided by the Moffitt Cancer Center’s Office of Scientific Publishing by Daley White and Gerard Hebert; no compensation was given beyond their regular salaries. This work has been supported in part by the Flow Cytometry, Small Animal Imaging Lab, and Tissue Core Facilities at the H. Lee Moffitt Cancer Center & Research Institute, an NCI designated Comprehensive Cancer Center (P30-CA076292). We gratefully acknowledge the funding provided by the Radiological Society of North America (RSNA) and express our appreciation to the Moffitt Vivarium staff for their invaluable assistance in the care of the animals used in this study. We also extend our thanks to Cellular Nanomed for supplying the magnetoelectric nanoparticles essential to this research. The segmentation work in this study was facilitated by 3D Slicer, a free and open-source software platform for the analysis and visualization of medical and biomedical images. We also acknowledge Chen-Yu Chang, a researcher apart of Professor Daniela Radu’s group at the Florida International University, who helped us obtain high-resolution TEM images of the nanoparticles.

## Author contributions

Conception or design of the work: J.M.B., V.A., M.S., K.H., J.W., I.O., V.K., K.Y., S.H., P.L., S.K., R.A.G., M.M. Acquisition of data: J.M.B., E.S., V.A., M.S., E.Z., V.E., K.H., D.M., A.S.L., D.A., S.C. (Shawnus Chen), M.A.M., S.C. (Skye Conlan), J.A.G., S.P.T., G.R., K.L., N.R., J.M.F. Data analysis: J.M.B., V.A., M.S., K.H., J.C., S.K. Interpretation of data: J.M.B., V.A., M.S., K.H., S.K., R.A.G., M.M., P.L. Drafting of the work: J.M.B. E.S., V.A., M.S., J.W., S.K.

## Materials & Correspondence

Correspondence and material requests should be addressed to John Michael Bryant, MD.

## Additional information

This article includes [7] Extended Data Tables and [4] Extended Data Figures.

## Data Availability Statement

The data that support the findings of this study are available from the corresponding author upon reasonable request.

