## Supplement for "Nanoparticles use magnetoelectricity to target and eradicate cancer cells"

**SUPPLEMENTARY MATERIAL**

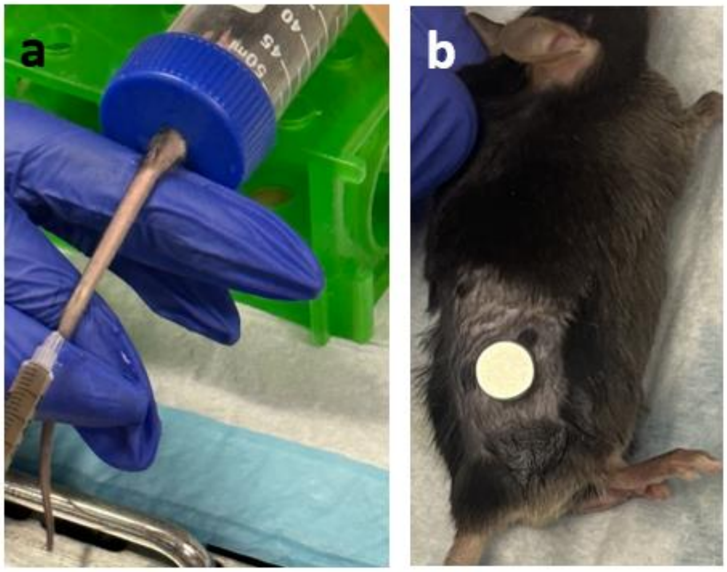


**Extended Data Fig. S1: Photo examples of tail vein injection and neodymium magnet placement for tumor targeting**. **a** Demonstration of a typical tail vein injection. The mice were placed within a 50 mL Eppendorf conical tube with a small hole placed through the top to pull the tail through. Every mouse in the study received a 300 mL tail vein injection in between the M0 and M1 timepoints. **b** demonstration of the neodymium magnet, generating a 0.15 T field over the flank tumors, (5.64 mm height by 4.0 mm radius) placement over the murine flank tumors. Adhesion was accomplished with a small application of VetBond Tissue Adhesive (3M, Saint Paul, MN, USA) to the flank tumor and subsequent placement of the neodymium magnet. These magnets were removed the next day immediately prior to MRI via gentle pulling when the mice were anesthetized.

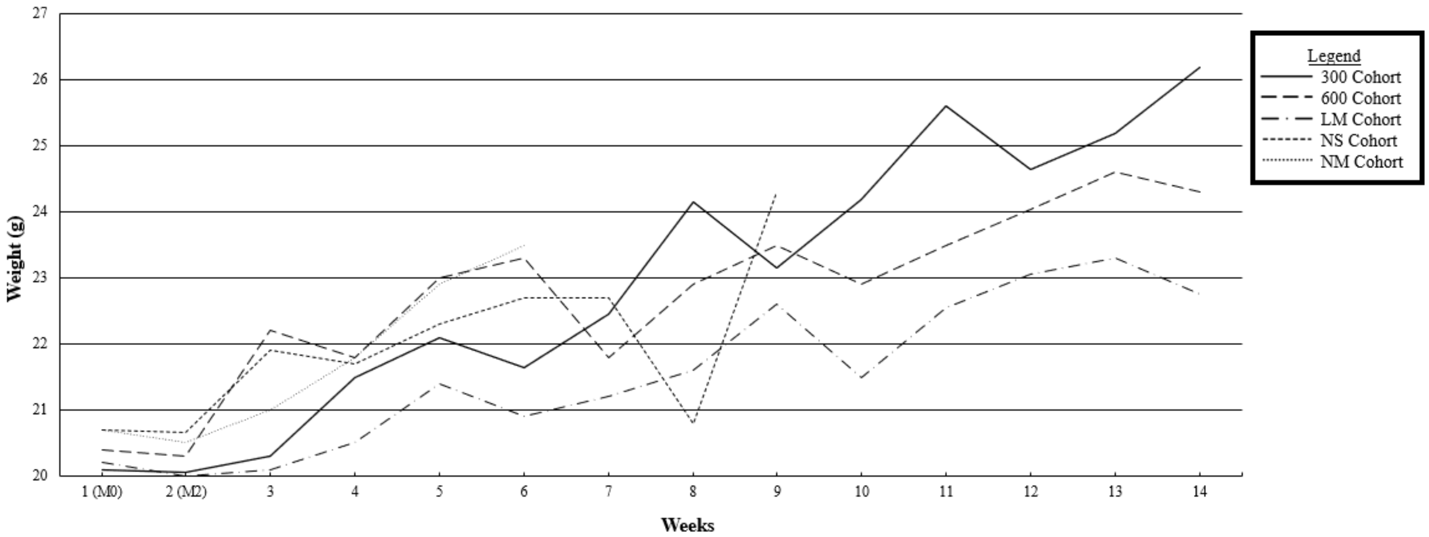


**Extended Data Fig. S2: Confirmatory study median per cohort mouse weights from M0 to end of study**. Line graph of median mouse weights per cohort from week 1 to week 14 of the confirmatory study. In general, the mice gained weight over the course of the study with mild-to-moderate drops as their tumor burden approached 1,000 $mm^3$.

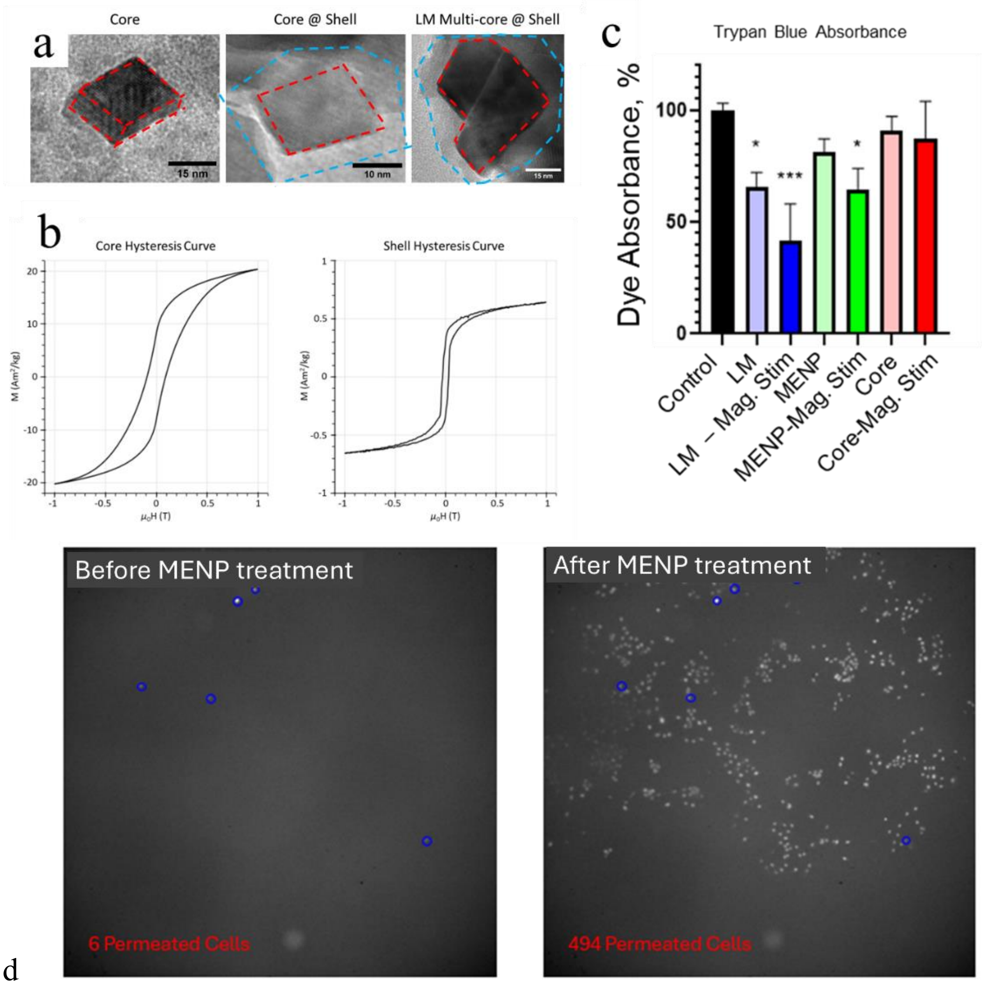


**Extended Data Fig. S3: MENPs characterization and ME measurements**. **a** HR-TEM Images of the magnetostrictive cobalt-ferrite-based inverse-spinel ferrimagnetic core of a rectangular prism shape and the core-shell nanostructure with the piezoelectric barium-titanite-based perovskite shell enclosing the magnetostrictive core for the default MENPs (e.g., as used within the 300 and 600 cohorts). The core and the shell are highlighted by red and blue broken lines, respectively. The multi-core-shell of the low-ME MENPs (LM cohort) is shown to the right. **b** M-H loops of the core nanoparticles and the core-shell nanoparticles, measured with AGM Lakeshore Nanomag 2900 (Lake Shore Cryotronics, Westerville, OH, USA). **c** Summary of the results of the Trypan Blue dye test optimization of the ME effect of MENPs[1]. Standard MENPs had no significant effect on the dye solution on their own, but upon stimulation with an AC magnetic field, the dye absorption was reduced to 41.9% of the original absorption ($p$=0.0002). This corresponds to an ME effect on the order of 1 V/A[2]. Similarly, the LMs only demonstrated an effect on dye absorption upon AC magnetic field stimulation, with a reduction in absorption of 64.8% (p=0.0115). This corresponds to a ME effect on the order of 0.5 V/A, which is equivalent to 150 µg of standard MENPs). MENP cores (i.e., naked ferrimagnetic cores) were tested as a control and validation of the test and did not have any effect on the dye absorption regardless of AC stimulation. **d** IRE verification in other in vitro studies. Microscopy images of SKOV-3 cells before and after MENP treatment using propidium iodide to determine membrane permeability. Blue outlines show cells permeated before and after treatment. As propidium iodide is membrane impermeable, but fluoresces upon reaction with nucleic acids, it is a useful marker of electroporation. This simple proof-of-concept experiment demonstrates the ability of MENPs to electroporate cancer cells in vitro. 4 ul of MENP solution was added to SKOV-3 cells plated in 2-ml McCoys 5A Media (Life Technologies, NY) and stimulated with a 30-minute magnetic field treatment of 0.1-T 20-ms pulses every 5 seconds.

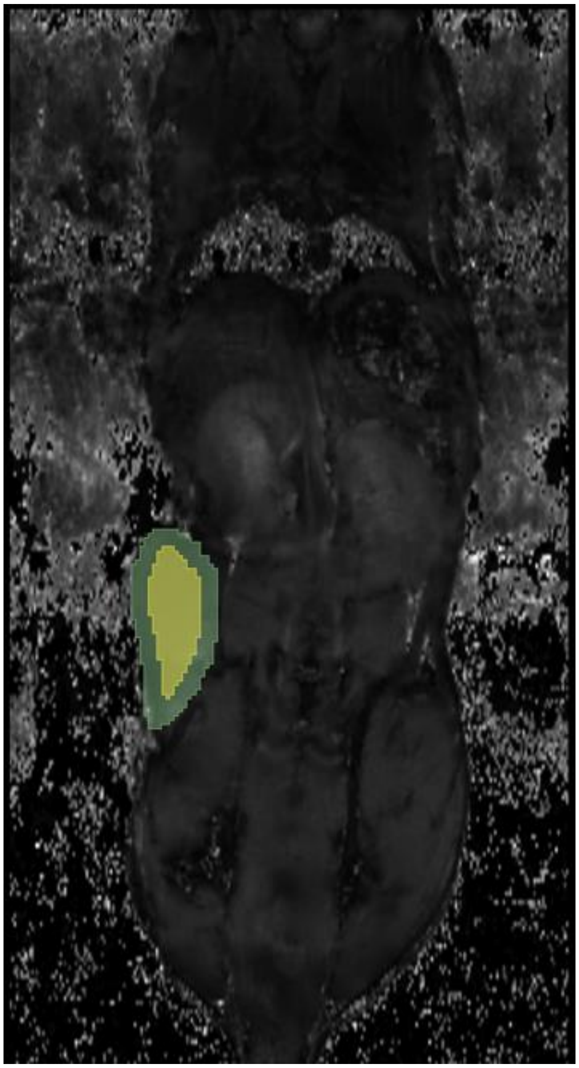

**Extended Data Fig. S4: Contouring example**. Coronal slice from an baseline (M0) $T_2$ map. Green segment represents volumetric contour and yellow segment represents a isotropic 1 mm contraction to control for subtle motion and volumetric averaging.

**Extended Data Table S1.** Pilot study median tumor volume, relaxometry, and weight data

| Metric | Cohort 60 (n=5) | Cohort 300 (n=5) | Cohort 600 (n=5) | Cohort LM (n=3) | Cohort NS (n=3) |
|---|---|---|---|---|---|
| M0 tumor volume, mm³ (IQR) | 87.7 (67.8-110.8) | 152.3 (119.5-155.8) | 53.4 (41.4-88.1) | 39.6 (32.6-48.9) | 41.2 (35.9-69.2) |
| M2 tumor volume, mm³ (IQR) | 97.9 (66.2-188.4) | 76.6 (60.0-126.7) | 26.6 (20.9-43.8) | 79.9 (63.7-85.9) | 81.1 (68.9-171.0) |
| M2/M0 tumor volume, % (IQR) | 174.0% (111.6%-237.6%) | 72.3% (50.3%-81.3%) | 49.8% (47.0%-49.8%) | 185.8% (161.6%-209.0%) | 265.0% (201.3%-266.8%) |
| M0 tumor T2*, ms (IQR) | 12.4 (10.3-13.8) | 10.7 (8.9-12.8) | 10.3 (9.2-11.3) | 11.0 (10.2-12.3) | 7.7 (7.0-9.1) |
| M1 tumor T2*, ms (IQR) | 12.0 (10.8-13.2) | 6.9 (5.2-7.5) | 6.1 (6.0-6.1) | 14.3 (13.8-15.1) | 12.3 (10.9-12.7) |
| M1/M0 tumor T2*, % (IQR) | 97.2% (96.9%-105.6%) | 61.4% (58.9%-64.2%) | 59.1% (58.1%-60.9%) | 122.2% (113.6%-145.8%) | 151.7% (139.3%-155.4%) |
| M0 tumor T2, ms (IQR) | 44.3 (43.7-50.1) | 57.3 (57.1-67.0) | 52.5 (49.1-57.7) | 49.0 (47.7-53.7) | 53.1 (52.4-54.0) |
| M1 tumor T2, ms (IQR) | 56.1 (50.9-68.0) | 51.1 (49.9-54.6) | 51.9 (36.7-54.0) | 57.0 (52.3-63.2) | 55.1 (54.7-55.6) |
| M1/M0 tumor T2, % (IQR) | 155.5% (101.6%-159.6%) | 86.2% (83.9%-87.4%) | 93.7% (60.5%-94.2%) | 97.7% (97.3%-123.4%) | 102.1% (101.2%-105.2%) |
| M0 mice weights, g (IQR) | 26.3 (24.9-26.5) | 25.3 (21.9-25.6) | 26.1 (20.8-26.1) | 20.2 (19.9-23.7) | 25.2 (22.8-26.3) |
| M2 mice weights, g (IQR) | 25.5 (24.4-25.7) | 25.5 (21.7-25.5) | 24.0 (19.8-25.4) | 20.0 (19.9-23.5) | 26.7 (23.7-26.9) |

***Abbreviations.*** 60: 60 µg of MENPs cohort; 300: 300 µg of MENPs cohort; 600: 600 µg of MENPs cohort; IQR: interquartile range; LM: 300 µg of low-ME coefficient nanoparticles cohort; M0: timepoint M0; M1: timepoint M1; M2: timepoint M2; NS: normal saline control cohort.

**Extended Data Table S2.** Fisher's least significant difference comparisons for the pilot study.

**Relative tumor volume at time M2 from time M0 (%)**

| (I) Cohort | (J) Cohort | Mean Difference | Std. Error | P value | 95% CI Lower Bound | 95% CI Upper Bound |
|---|---|---|---|---|---|---|
| 60ug | 300ug | 106.3%* | 35.3% | **.008** | 31.4% | 181.2% |
| | 600ug | 124.2%* | 35.3% | **.003** | 49.3% | 199.1% |
| | LM | -13.0% | 40.8% | .753 | -99.5% | 73.4% |
| | NS | -51.6% | 40.8% | .224 | -138.1% | 34.9% |
| 300ug | 600ug | 17.9% | 35.3% | .619 | -57.0% | 92.8% |
| | LM | -119.3%* | 40.8% | **.010** | -205.8% | -32.8% |
| | NS | -157.9%* | 40.8% | **.001** | -244.4% | -71.4% |

| | | | | | | |
|---|---|---|---|---|---|---|
| 600ug | LM | -137.3%* | 40.8% | **.004** | -223.7% | -50.8% |
| | NS | -175.8%* | 40.8% | **<.001** | -262.3% | -89.4% |
| LM | NS | -38.6% | 45.6% | .410 | -135.3% | 58.1% |
| **Absolute tumor volume differences at time M2 from time M0 ($mm^3$)** | | | | | | |
| (I) Cohort | (J) Cohort | Mean Difference | Std. Error | P value | 95% CI Lower Bound | 95% CI Upper Bound |
| 60ug | 300ug | 122.1 * | 47.2 | **.020** | 21.9 | 222.3 |
| | 600ug | 118* | 47.2 | **.024** | 17.8 | 218.1 |
| | LM | 44.3 | 54.6 | .428 | -71.3 | 160.0 |
| | NS | -.2 | 54.6 | .997 | -115.9 | 115.4 |
| 300ug | 600ug | -4.1 | 47.2 | .931 | -104.3 | 96.0 |
| | LM | -77.8 | 54.6 | .173 | -193.4 | 37.9 |
| | NS | -122.3* | 54.6 | **.039** | -238.0 | -6.7 |
| 600ug | LM | -73.6 | 54.6 | .196 | -189.3 | 42.0 |
| | NS | -118.2 * | 54.6 | **.046** | -233.8 | -2.5 |
| LM | NS | -44.6 | 61.0 | .476 | -173.9 | 84.7 |
| **Relative $T_2^*$ relaxation time at time M1 from time M0 (%)** | | | | | | |
| (I) Cohort | (J) Cohort | Mean Difference | Std. Error | P value | 95% CI Lower Bound | 95% CI Upper Bound |
| 60ug | 300ug | 76.9* | 31.5% | **.026** | 10.5% | 143.4% |
| | 600ug | 55.9% | 31.4% | .094 | -10.6% | 122.4% |
| | LM | -19.8% | 36.2% | .592 | -96.6% | 57.0% |
| | NS | -4.3% | 36.2% | .907 | -81.1% | 72.5% |
| 300ug | 600ug | -21.0% | 31.4% | .512 | -87.5% | 45.4% |
| | LM | -96.7%* | 36.2% | **.017** | -173.5% | -20.0% |
| | NS | -81.2%* | 36.2% | **.039** | -158.0% | -4.5% |
| 600ug | LM | -75.7% | 36.2% | .053 | -152.5% | 1.1% |
| | NS | -60.2% | 36.2% | .116 | -137.0% | 16.6% |
| LM | NS | 15.5% | 40.5% | .707 | -70.3% | 101.3% |
| **Absolute $T_2^*$ relaxation time differences at time M1 from time M0 (ms)** | | | | | | |
| (I) Cohort | (J) Cohort | Mean Difference | Std. Error | P value | 95% CI Lower Bound | 95% CI Upper Bound |
| 60ug | 300ug | 7.8* | 2.8 | **.014** | 1.8 | 13.7 |
| | 600ug | 3.9 | 2.8 | .187 | -2.1 | 9.8 |
| | LM | .3 | 3.2 | .932 | -6.6 | 7.2 |
| | NS | -1.0 | 3.2 | .767 | -7.8 | 5.9 |
| 300ug | 600ug | -3.9 | 2.8 | .182 | -9.9 | 2.0 |
| | LM | -7.5* | 3.2 | **.034** | -14.4 | -.6 |
| | NS | -8.8 * | 3.2 | **.016** | -15.6 | -1.9 |
| 600ug | LM | -3.6 | 3.2 | .284 | -10.5 | 3.3 |
| | NS | -4.8 | 3.2 | .154 | -11.7 | 2.0 |
| LM | NS | -1.3 | 3.6 | .733 | -8.9 | 6.4 |

| **Relative $T_2$ relaxation time at time M1 from time M0 (%)** | | | | | | |
|---|---|---|---|---|---|---|
| (I) Cohort | (J) Cohort | Mean Difference | Std. Error | P value | 95% CI Lower Bound | 95% CI Upper Bound |
| 60ug | 300ug | 52.9%* | 16.8% | **.006** | 17.2% | 88.5% |
| | 600ug | 59.5%* | 16.8% | **.003** | 23.9% | 95.2% |
| | LM | 23.3% | 19.4% | .247 | -17.9% | 64.5% |
| | NS | 34.4% | 19.4% | .096 | -6.8% | 75.5% |
| 300ug | 600ug | 6.6% | 16.8% | .699 | -29.0% | 42.3% |
| | LM | -29.6% | 19.4% | .147 | -70.7% | 11.6% |
| | NS | -18.5% | 19.4% | .354 | -59.7% | 22.6% |
| 600ug | LM | -36.2% | 19.4% | .081 | -77.4% | 5.0% |
| | NS | -25.1% | 19.4% | .214 | -66.3% | 16.0% |
| LM | NS | 11.1% | 21.7% | .618 | -35.0% | 57.1% |
| **Absolute $T_2$ relaxation time differences at time M1 from time M0 (ms)** | | | | | | |
| (I) Cohort | (J) Cohort | Mean Difference | Std. Error | P value | 95% CI Lower Bound | 95% CI Upper Bound |
| 60ug | 300ug | 24.5* | 9.3 | **.018** | 4.9 | 44.1 |
| | 600ug | 30.1* | 9.3 | **.005** | 10.5 | 49.7 |
| | LM | 8.7 | 10.7 | .430 | -14.0 | 31.3 |
| | NS | 13.5 | 10.7 | .226 | -9.2 | 36.1 |
| 300ug | 600ug | 5.6 | 9.3 | .554 | -14.0 | 25.2 |
| | LM | -15.8 | 10.7 | .158 | -38.5 | 6.8 |
| | NS | -11.0 | 10.7 | .318 | -33.7 | 11.6 |
| 600ug | LM | -21.4 | 10.7 | .062 | -44.1 | 1.2 |
| | NS | -16.6 | 10.7 | .140 | -39.3 | 6.0 |
| LM | NS | 4.8 | 12.0 | .692 | -20.5 | 30.2 |
| Abbreviations: CI: Confidence Interval; LM: low magnetoelectric effect MENPs; NS: normal saline | | | | | | |

* The mean difference is significant at the 0.05 level.

Bolded P values are significant at 0.05 level

| **Extended Data Table S3.** Confirmatory study median tumor volume, relaxometry, and survival data | | | | | |
|---|---|---|---|---|---|
| **Metric** | **Cohort 300 (n=6)** | **Cohort 600 (n=6)** | **Cohort LM (n=5)** | **Cohort NS (n=5†)** | **Cohort NM (n=5)** |
| M0 tumor volume, $mm^3$ (IQR) | 137.1 (112.9-241.1) | 108.0 (49.2-227.9) | 83.7 (20.6-212.7) | 107.5 (80.3-128.0) | 84.9 (38.2-199.1) |
| M2 tumor volume, $mm^3$ (IQR) | 97.5 (45.5-131.3) | 86.5 (23.3-223.4) | 78.8 (25.9-316.6) | 166.0 (119.8-242.8) | 140.3 (66.1-463.7) |
| M2/M0 tumor volume, % (IQR) | 56.7% (49.2%-81.5%) | 73.7% (46.9%-98.6%) | 148.8% (125.6%-150.5%) | 159.4% (156.8%-180.5%) | 163.4% (148.3%-202.6%) |

| | | | | | |
|---|---|---|---|---|---|
| M0 tumor $T_2$*, ms (IQR) | 11.2 (9.5-13.6) | 10.1 (8.9-14.4) | 9.5 (8.4-9.7) | 12.8 (12.0-13.0) | NA |
| M1 tumor $T_2$*, ms (IQR) | 9.9 (9.4-11.6) | 8.0 (5.9-9.0) | 9.7 (9.6-12.9) | 12.5 (10.2-14.3) | NA |
| M1/M0 tumor $T_2$*, % (IQR) | 88.8% (82.2%-96.5%) | 86.1% (72.2%-88.5%) | 132.3% (114.5%-132.4%) | 110.1% (104.6%-116.8%) | NA |
| M0 tumor $T_2$, ms (IQR) | 43.5 (35.9-44.2) | 46.9 (42.8-58.5) | 45.6 (43.0-49.2) | 37.6 (35.8-40.2) | NA |
| M1 tumor $T_2$, ms (IQR) | 39.3 (35.9-41.9) | 44.7 (42.4-46.0) | 47.3 (46.5-48.9) | 38.2 (38.1-38.9) | NA |
| M1/M0 tumor $T_2$, % (IQR) | 97.4% (81.2%-103.7%) | 85.8% (75.9%-101.5%) | 102.0% (96.1%-115.2%) | 110.3% (101.0%-102.7%) | NA |
| Survival time from M1, days (IQR) | 31.5 (27.0-81.8) | 40.8 (28.8-88.3) | 33.0 (12.0-97.0) | 33.0 (12.0-33.0) | 33.0 (13.0-33.0) |
| Number to Reach Tumor Volume Endpoint* | 1 | 3 | 1 | 0 | 2 |
| Number to Reach Tumor Diameter Endpoint** | 1 | 0 | 1 | 1 | 2 |
| Number to Reach Ulcer Diameter Endpoint*** | 2 | 1 | 1 | 4 | 1 |
| Number to reach CR | 2 | 2 | 2 | 0 | 0 |
| Time to CR from M1, days (IQR) | 17.0 (15.5-18.5) | 14.0 (14.0-14.0) | 30.5 (28.8-32.3) | NA | NA |

*tumor volume ≥1,000 $mm^3$
**tumor diameter ≥1 cm
***tumor ulcer diameter ≥5 mm
†Tumor volume measurements only include four mice as one mouse within NS cohort reached endpoint after M1 but before M2.
***Abbreviations.*** 300: 300 µg of MENPs cohort; 600: 600 µg of MENPs cohort; IQR: interquartile range; LM: 300 µg of low-ME MENPs cohort; NA: not applicable; NM: no M1 timepoint magnetic resonance imaging control cohort; NS: normal saline control cohort.

| **Extended Data Table S4.** Fisher's least significant difference comparisons for the confirmatory study. |
|---|
| **Relative tumor volume at time M2 from time M0 (%)** |

| (I) Cohort | (J) Cohort | Mean Difference | Std. Error | P value | 95% CI Lower Bound | 95% CI Upper Bound |
|---|---|---|---|---|---|---|
| 300ug | 600ug | -9.7% | 20.3% | .638 | -51.9% | 32.5% |
| | LM | -82.0%* | 21.3% | **<.001** | -126.3% | -37.8% |
| | NM | -114.2%* | 21.3% | **<.001** | -158.5% | -70.0% |
| | NS | -115.9%* | 22.7% | **<.001** | -163.1% | -68.7% |
| 600ug | LM | -72.3%* | 21.3% | **.003** | -116.6% | -28.1% |
| | NM | -104.6%* | 21.3% | **<.001** | -148.8% | -60.3% |
| | NS | -106.2%* | 22.7% | **<.001** | -153.4% | -59.0% |
| LM | NM | -32.2% | 22.2% | .162 | -78.5% | 14.0% |
| | NS | -33.9% | 23.6% | .166 | -82.9% | 15.2% |
| NM | NS | -1.7% | 23.6% | .944 | -50.7% | 47.4% |
| **Absolute tumor volume differences at time M2 from time M0 ($mm^3$)** | | | | | | |
| (I) Cohort | (J) Cohort | Mean Difference | Std. Error | P value | 95% CI Lower Bound | 95% CI Upper Bound |
| 300ug | 600ug | -43.3 | 55.3 | .443 | -158.3 | 71.8 |
| | LM | -140.6 * | 58.0 | **.024** | -261.2 | -20.0 |
| | NM | -189.7* | 58.0 | **.004** | -310.3 | -69.1 |
| | NS | -154.5 * | 61.8 | **.021** | -283.1 | -25.9 |
| 600ug | LM | -97.3 | 58.0 | .108 | -218.0 | 23.3 |
| | NM | -146.4* | 58.0 | **.020** | -267.1 | -25.8 |
| | NS | -111.2 | 61.8 | .086 | -239.8 | 17.4 |
| LM | NM | -49.1 | 60.6 | .427 | -175.1 | 76.9 |
| | NS | -13.9 | 64.3 | .831 | -147.5 | 119.8 |
| NM | NS | 35.2 | 64.3 | .589 | -98.4 | 168.9 |
| **Relative $T_2^*$ relaxation time at time M1 from time M0 (%)** | | | | | | |
| (I) Cohort | (J) Cohort | Mean Difference | Std. Error | P value | 95% CI Lower Bound | 95% CI Upper Bound |
| 300ug | 600ug | 2.3% | 11.6% | .845 | -22.1% | 26.8% |
| | LM | -32.1%* | 12.2% | **.017** | -57.8% | -6.5% |
| | NS | -21.3 % | 12.2% | .098 | -46.9% | 4.4% |
| 600ug | LM | -34.5 %* | 12.2% | **.011** | -60.1% | -8.8% |
| | NS | -23.6 % | 12.2% | .069 | -49.2% | 2.0% |
| LM | NS | 10.9 % | 12.7% | .406 | -15.9% | 37.6% |
| **Absolute $T_{2*}$ relaxation time differences at time M1 from time M0 (ms)** | | | | | | |
| (I) Cohort | (J) Cohort | Mean Difference | Std. Error | P value | 95% CI Lower Bound | 95% CI Upper Bound |
| 300ug | 600ug | .1 | 1.1 | .909 | -2.1 | 2.3 |
| | LM | -2.9* | 1.1 | **.018** | -5.2 | -.6 |
| | NS | -2.5* | 1.1 | **.034** | -4.9 | -.2 |
| 600ug | LM | -3.0* | 1.1 | **.014** | -5.4 | -.7 |
| | NS | -2.7 * | 1.1 | **.027** | -5.0 | -.3 |

| | | | | | | |
|---|---|---|---|---|---|---|
| LM | NS | .4 | 1.2 | .764 | -2.1 | 2.8 |
| **Relative $T_2$ relaxation time at time M1 from time M0 (%)** | | | | | | |
| (I) Cohort | (J) Cohort | Mean Difference | Std. Error | P value | 95% CI Lower Bound | 95% CI Upper Bound |
| 300ug | 600ug | 7.9% | 10.2% | .452 | -13.6% | 29.4% |
| | LM | -17.1% | 10.7% | .129 | -39.6% | 5.5% |
| | NS | -7.9% | 10.7% | .471 | -30.5% | 14.7% |
| 600ug | LM | -25.0%* | 10.7% | **.032** | -47.5% | -2.4% |
| | NS | -15.8% | 10.7% | .159 | -38.4% | 6.8% |
| LM | NS | 9.2% | 11.2% | .424 | -14.4% | 32.7% |
| **Absolute $T_2$ relaxation time differences at time M1 from time M0 (ms)** | | | | | | |
| (I) Cohort | (J) Cohort | Mean Difference | Std. Error | P value | 95% CI Lower Bound | 95% CI Upper Bound |
| 300ug | 600ug | 9.2 | 7.4 | .230 | -6.4 | 24.8 |
| | LM | -8.0 | 7.8 | .318 | -24.3 | 8.3 |
| | NS | -4.0 | 7.8 | .609 | -20.4 | 12.3 |
| 600ug | LM | -17.2 * | 7.8 | **.040** | -33.5 | -.9 |
| | NS | -13.2 | 7.8 | .105 | -29.6 | 3.1 |
| LM | NS | 3.9 | 8.1 | .633 | -13.1 | 21 |
| Abbreviations: CI: Confidence Interval; LM: low magnetoelectric effect MENPs; NS: normal saline; NM: no magnetic field application<br>* The mean difference is significant at the 0.05 level.<br>Bolded P values are significant at 0.05 level | | | | | | |

**Extended Data Table S5**. Confirmatory study tissue collection and histological findings

| Cohort | **300** | **600** | **LM** | **NS** | **NM** |
|---|---|---|---|---|---|
| Number of mice in cohort | 6 | 6 | 5 | 5 | 5 |
| Number of mice with tissues collected | 5 | 5 | 5 | 4 | 3 |
| Bone Marrow (from femur) | | | | | |
| Hypocellular | 0 | 0 | 0 | 0 | 0 |
| Necrosis | 0 | 0 | 0 | 0 | 0 |
| Fibrosis | 0 | 0 | 0 | 0 | 0 |
| Stomach & Duodenum | | | | | |
| Focal chronic inflammation | 5 (100%) | 5 (100%) | 5 (100%) | 4 (100%) | 3 (100%) |
| Epithelial degeneration | 0 | 0 | 0 | 0 | 0 |
| Epithelial hyperplasia | 0 | 0 | 0 | 0 | 0 |
| Necrosis | 0 | 0 | 0 | 0 | 0 |
| Fibrosis | 0 | 0 | 0 | 0 | 0 |
| Heart | | | | | |
| Myocardial vacuolation | 0 | 0 | 0 | 0 | 0 |
| Hemorrhage | 0 | 0 | 0 | 0 | 0 |
| Inflammation (myocardium/valve) | 0 | 0 | 0 | 0 | 0 |
| Necrosis | 0 | 0 | 0 | 0 | 0 |
| Fibrosis | 0 | 0 | 0 | 0 | 0 |

| | | | | | |
|---|---|---|---|---|---|
| Kidneys | | | | | |
| Diffuse tubule necrosis, mild | 5 (100%) | 2 (40.0%) | 4 (80.0%) | 2 (50.0%) | 2 (66.7%) |
| Diffuse tubule necrosis, moderate/severe | 0 | 0 | 0 | 0 | 0 |
| Glomerulosclerosis | 0 | 0 | 0 | 0 | 0 |
| Tubule hyperplasia | 0 | 0 | 0 | 0 | 0 |
| Interstitial inflammation, chronic | 0 | 2 (40.0%) | 0 | 1 (25.0%) | 0 |
| Liver | | | | | |
| Extramedullary hematopoiesis, mild | 5 (100%) | 5 (100%) | 5 (100%) | 4 (100%) | 3 (100%) |
| Extramedullary hematopoiesis, moderate/severe | 0 | 0 | 0 | 0 | 0 |
| Cytoplasmic vacuolation consistent with glycogen | 1 (20.0%) | 3 (60.0%) | 1 (20.0%) | 0 | 0 |
| Centrilobular necrosis | 0 | 0 | 0 | 0 | 0 |
| Centrilobular fibrosis | 0 | 0 | 0 | 0 | 0 |
| Kupffer cell hyperplasia | 0 | 0 | 0 | 0 | 0 |
| Lungs, | | | | | |
| Alveolar/bronchiolar epithelial hyperplasia, mild | 2 (40.0%) | 3 (60.0%) | 4 (80.0%) | 4 (100%) | 3 (100%) |
| Alveolar/bronchiolar epithelial hyperplasia, | 0 | 0 | 0 | 0 | 0 |
| moderate/severe | 5 (100%) | 3 (60.0%) | 5 (100%) | 4 (100%) | 3 (100%) |
| Vascular congestion, moderate | 0 | 0 | 0 | 0 | 0 |
| Vascular congestion, severe | 0 | 0 | 0 | 0 | 0 |
| Histocyte cellular infiltration | 0 | 0 | 0 | 0 | 0 |
| Focal necrosis | 0 | 0 | 0 | 0 | 0 |
| Granulomatous chronic inflammation | 0 | 0 | 0 | 2 (50.0%) | 1 (33.3%) |
| Focal chronic inflammation | 0 | 1 (20.0%) | 0 | 2 (50.0%) | 1 (33.3%) |
| Septal fibrosis, mild | 0 | 0 | 0 | 0 | 0 |
| Septal fibrosis, moderate/severe | | | | | |
| Spleen | | | | | |
| Extramedullary hematopoiesis, mild | 0 | 0 | 2 (40.0%) | 1 (25.0%) | 0 |
| Extramedullary hematopoiesis, moderate | 3 (60.0%) | 4 (80.0%) | 2 (40.0%) | 1 (25.0% | 1 (33.3%) |
| Extramedullary hematopoiesis, severe | 2 (40.0%) | 1 (20.0%) | 1 (20.0%) | 2 (50.0%) | 2 (66.7%) |
| Reactive chronic inflammation, mild | 0 | 0 | 1 (20.0%) | 0 | 0 |
| Reactive chronic inflammation, moderate | 2 (40.0%) | 3 (60.0%) | 3 (60.0%) | 3 (75.0%) | 3 (100%) |
| Reactive chronic inflammation, severe | 3 (60.0%) | 2 (40.0%) | 1 (20.0%) | 1 (25.0%) | 0 |
| Vascular congestion, mild | 0 | 0 | 0 | 0 | 0 |
| Vascular congestion, moderate | 5 (100%) | 5 (100%) | 5 (100%) | 4 (100%) | 3 (100%) |
| Fibrosis | 0 | 0 | 0 | 0 | 0 |

***Abbreviations.*** 300: 300 µg of magnetoelectric nanoparticles cohort; 600: 600 µg of magnetoelectric nanoparticles cohort; LM: 300 µg of low-magnetoelectric coefficient nanoparticle cohort; NM: no M1 timepoint magnetic resonance imaging control cohort; NS: normal saline control cohort.

**Extended Data Table S6.** Combined pilot and confirmatory study tumor volume and relaxometry data

| Metric | Cohort 300 (n=11) | Cohort 600 (n=11) | Cohort LM (n=8) | Cohort NS (n=8†) | Cohort 60 (n=5) | Cohort NM (n=5) |
|---|---|---|---|---|---|---|
| M0 tumor volume, $mm^3$ (IQR) | 146.3 (113.7-214.2) | 81.9 (39.8-153.5) | 48.9 (24.4-116.0) | 97.1 (35.9-107.5) | 87.7 (67.8-110.8) | 84.9 (38.2-199.1) |

| | | | | | | |
|---|---|---|---|---|---|---|
| M2 tumor volume, mm$^3$ (IQR) | 79.0 (48.5-131.5) | 40.5 (19.2-102.9) | 79.3 (42.2-148.1) | 157.3 (68.9-217.8) | 97.9 (66.2-188.4) | 140.3 (66.1-463.7) |
| M2/M0 tumor volume, % (IQR) | 59.3% (49.0%-85.1%) | 49.8% (46.5%-74.2%) | 149.6% (134.5%-189.7%) | 159.4% (154.3%-254.4%) | 174.0% (111.6%-237.6%) | 159.4% (156.8%-180.5%) |
| M0 tumor $T_{2*}$, ms (IQR) | 10.9 (8.9-13.3) | 10.3 (9.0-11.5) | 9.6 (9.2-11.6) | 11.2 (7.6-12.9) | 12.4 (10.3-13.8) | NA |
| M1 tumor $T_{2*}$, ms (IQR) | 8.4 (7.0-9.9) | 6.1 (5.8-8.0) | 13.2 (9.6-14.7) | 12.4 (10.0-13.5) | 12.0 (10.8-13.2) | NA |
| M1/M0 tumor $T_{2*}$, % (IQR) | 77.4% (62.8%-88.8%) | 67.9% (58.6%-86.1%) | 127.3% (112.1%-132.7%) | 121.8% (108.8%-134.6%) | 97.2% (96.9%-105.6%) | NA |
| M0 tumor $T_2$, ms (IQR) | 52.9 (43.4-57.2) | 49.1 (45.9-59.9) | 47.7 (45.0-49.5) | 41.4 (37.1-52.1) | 44.3 (43.7-50.1) | NA |
| M1 tumor $T_2$, ms (IQR) | 42.4 (39.3-50.5) | 45.5 (40.2-52.9) | 48.2 (47.1-58.5) | 41.0 (38.1-54.5) | 56.1 (50.9-68.0) | NA |
| M1/M0 tumor $T_2$, % (IQR) | 87.4% (81.2%-97.4%) | 91.2% (67.4%-97.4%) | 99.8% (96.8%-123.1%) | 101.7% (100.8%-104.1%) | 155.5% (101.6%-159.6%) | NA |

†tumor volume measurements only include four mice as one mouse within NS cohort reached endpoint after M1 but before M2.

***Abbreviations.*** 60: 60 µg of MENPs cohort; 300: 300 µg of MENPs cohort; 600: 600 µg of MENPs cohort; IQR: interquartile range; LM: 300 µg of low-ME coefficient nanoparticles cohort; M0: timepoint M0; M1: timepoint M1; M2: timepoint M2; NA: not applicable; NM: no M1 timepoint magnetic resonance imaging control cohort; NS: normal saline control cohort.

**Extended Data Table S7.** Fisher's least significant difference comparisons for both studies combined.

**Relative tumor volume at time M2 from time M0 (%)**

| (I) Cohort | (J) Cohort | Mean Difference | Std. Error | P value | 95% CI Lower Bound | 95% CI Upper Bound |
|---|---|---|---|---|---|---|

| | | | | | | |
|---|---|---|---|---|---|---|
| 60ug | 300ug | 108.3%* | 24.4% | **<.001** | 59.0% | 157.7% |
| | 600ug | 111.2%* | 24.4% | **<.001** | 61.8% | 160.6% |
| | LM | 12.6% | 25.8% | .628 | -39.6% | 64.8% |
| | NM | -4.2% | 28.7% | .884 | -62.1% | 53.7% |
| | NS | -25.5% | 26.5% | .343 | -79.1% | 28.1% |
| 300ug | 600ug | 2.9% | 19.3% | .883 | -36.2% | 41.9% |
| | LM | -95.7%* | 21.1% | **<.001** | -138.3% | -53.2% |
| | NM | -112.5%* | 24.4% | **<.001** | -161.9% | -63.2% |
| | NS | -133.8%* | 21.9% | **<.001** | -178.1% | -89.6% |
| 600ug | LM | -98.6%* | 21.1% | **<.001** | -141.1% | -56.1% |
| | NM | -115.4%* | 24.4% | **<.001** | -164.8% | -66.0% |
| | NS | -136.7%* | 21.9% | **<.001** | -180.9% | -92.4% |
| LM | NM | -16.8% | 25.8% | .519 | -69.0% | 35.4% |
| | NS | -38.1% | 23.5% | .112 | -85.5% | 9.3% |
| NM | NS | -21.3% | 26.5% | .427 | -74.9% | 32.3% |
| **Absolute tumor volume differences at time M2 from time M0 ($mm^3$)** | | | | | | |
| (I) Cohort | (J) Cohort | Mean Difference | Std. Error | P value | 95% CI Lower Bound | 95% CI Upper Bound |
| 60ug | 300ug | 129.1* | 45.3 | **.007** | 37.6 | 220.7 |
| | 600ug | 103.6* | 45.3 | **.027** | 12.1 | 195.2 |
| | LM | 13.1 | 47.9 | .786 | -83.7 | 109.9 |
| | NM | -54.7 | 53.2 | .309 | -162.1 | 52.6 |
| | NS | -11.2 | 49.2 | .820 | -110.6 | 88.1 |
| 300ug | 600ug | -25.5 | 35.8 | .481 | -97.9 | 46.9 |
| | LM | -116.0* | 39.1 | **.005** | -194.9 | -37.1 |
| | NM | -183.9 * | 45.3 | **<.001** | -275.4 | -92.3 |
| | NS | -140.4 * | 40.6 | **.001** | -222.4 | -58.3 |
| 600ug | LM | -90.5* | 39.1 | **.026** | -169.4 | -11.7 |
| | NM | -158.4 * | 45.3 | **.001** | -249.9 | -66.8 |
| | NS | -114.9 * | 40.6 | **.007** | -197.0 | -32.8 |
| LM | NM | -67.8 | 47.9 | .164 | -164.6 | 28.9 |
| | NS | -24.4 | 43.5 | .579 | -112.2 | 63.5 |
| NM | NS | 43.5 | 49.2 | .382 | -55.9 | 142.9 |
| **Relative $T_{2*}$ relaxation time at time M1 from time M0 (%)** | | | | | | |
| (I) Cohort | (J) Cohort | Mean Difference | Std. Error | P value | 95% CI Lower Bound | 95% CI Upper Bound |
| 60ug | 300ug | 59.2%* | 20.2 % | **.006** | 18.4% | 100.0% |
| | 600ug | 50.9%* | 20.2 % | **.016** | 10.1% | 91.7% |
| | LM | 0.3 % | 21.3% | .990 | -42.9% | 43.4% |

| | | | | | | |
|---|---|---|---|---|---|---|
| | NS | 12.9 % | 21.3% | .550 | -30.3% | 56.0% |
| 300ug | 600ug | -8.3% | 15.9% | .605 | -40.6% | 24.0% |
| | LM | -59.0%* | 17.4 % | **.002** | -94.1% | -23.8% |
| | NS | -46.4 %* | 17.4 % | **.011** | -81.5% | -11.2% |
| 600ug | LM | -50.7 %* | 17.4 % | **.006** | -85.8% | -15.5% |
| | NS | -38.1 %* | 17.4 % | **.035** | -73.2% | -2.9% |
| LM | NS | 12.6 % | 18.7 % | .504 | -25.2% | 50.4% |
| **Absolute $T_2^*$ relaxation time differences at time M1 from time M0 (ms)** | | | | | | |
| (I) Cohort | (J) Cohort | Mean Difference | Std. Error | P value | 95% CI Lower Bound | 95% CI Upper Bound |
| 60ug | 300ug | 5.2 * | 1.8 | **.008** | 1.5 | 8.9 |
| | 600ug | 3.5 | 1.8 | .067 | -.3 | 7.2 |
| | LM | .2 | 1.9 | .927 | -3.8 | 4.1 |
| | NS | -.01 | 1.9 | .972 | -4.0 | 3.9 |
| 300ug | 600ug | -1.7 | 1.5 | .247 | -4.7 | 1.2 |
| | LM | -5.0* | 1.6 | **.003** | -8.2 | -1.8 |
| | NS | -5.3 * | 1.6 | **.002** | -8.5 | -2.0 |
| 600ug | LM | -3.3 * | 1.6 | **.045** | -6.5 | -.1 |
| | NS | -3.5* | 1.6 | **.031** | -6.8 | -.3 |
| LM | NS | -.3 | 1.7 | .884 | -3.7 | 3.2 |
| **Relative $T_2$ relaxation time at time M1 from time M0 (%)** | | | | | | |
| (I) Cohort | (J) Cohort | Mean Difference | Std. Error | P value | 95% CI Lower Bound | 95% CI Upper Bound |
| 60ug | 300ug | 48.1%* | 11.5% | **<.001** | 24.8% | 71.4% |
| | 400ug | 55.4%* | 11.5% | **<.001** | 32.1% | 78.8% |
| | LM | 25.6%* | 12.2% | **.042** | 1.0% | 50.3% |
| | NS | 35.5%* | 12.2% | **.006** | 10.8% | 60.2% |
| 300ug | 600ug | 7.3% | 9.1% | .428 | -11.1% | 25.8% |
| | LM | -22.5%* | 9.9% | **.030** | -42.6% | -2.4% |
| | NS | -12.6% | 9.9% | .213 | -32.7% | 7.5% |
| 600ug | LM | -29.8%* | 9.9% | **.005** | -49.9% | -9.7% |
| | NS | -19.9% | 9.9% | .052 | -40.0% | 0.2% |
| LM | NS | 9.9% | 10.7% | .361 | -11.8% | 31.5% |
| **Absolute $T_2$ relaxation time differences at time M1 from time M0 (ms)** | | | | | | |
| (I) Cohort | (J) Cohort | Mean Difference | Std. Error | P value | 95% CI Lower Bound | 95% CI Upper Bound |
| 60ug | 300ug | 21.4 * | 7.1 | **.004** | 7.1 | 35.7 |
| | 600ug | 29.0* | 7.1 | **<.001** | 14.7 | 43.2 |
| | LM | 10.0 | 7.5 | .188 | -5.1 | 25.1 |

| | | | | | | |
|---|---|---|---|---|---|---|
| | NS | 14.3 | 7.5 | .063 | -.8 | 29.4 |
| 300ug | 600ug | 7.6 | 5.6 | .183 | -3.7 | 18.9 |
| | LM | -11.4 | 6.1 | .069 | -23.7 | .9 |
| | NS | -7.1 | 6.1 | .249 | -19.4 | 5.2 |
| 600ug | LM | -18.9* | 6.1 | **.003** | -31.3 | -6.6 |
| | NS | -14.7 * | 6.1 | **.021** | -27.0 | -2.4 |
| LM | NS | 4.3 | 6.5 | .518 | -9 | 17.5 |

Abbreviations: CI: Confidence Interval; LM: low magnetoelectric effect MENPs; NS: normal saline; NM: no magnetic field application
* The mean difference is significant at the 0.05 level.
Bolded P values are significant at 0.05 level


Extended Data References

1 Mushtaq, F. *et al.* Magnetoelectrically Driven Catalytic Degradation of Organics. *Adv Mater* **31**, e1901378 (2019). https://doi.org:10.1002/adma.201901378

2 Wang, P. *et al.* Colossal Magnetoelectric Effect in Core-Shell Magnetoelectric Nanoparticles. *Nano Lett* **20**, 5765-5772 (2020). https://doi.org:10.1021/acs.nanolett.0c01588
